# L1 retrotransposition is regulated post-transcriptionally In High-Grade Serous Ovarian Cancer

**DOI:** 10.1101/2022.09.27.509826

**Authors:** Barun Pradhan, Kaiyang Zhang, Yilin Li, Kari Lavikka, Taru Muranen, Kaisa Huhtinen, Richard Badge, Kathleen H. Burns, Johanna Hynninen, Sakari Hietanen, Jaana Oikkonen, Sampsa Hautaniemi, Liisa Kauppi

**Affiliations:** Research Program in Systems Oncology, Research Programs Unit, Faculty of Medicine, University of Helsinki, Helsinki, Finland; Cancer Research Unit, Institute of Biomedicine and FICAN West Cancer Centre, University of Turku, Turku, Finland; Department of Genetics and Genome Biology, University of Leicester, Leicester, United Kingdom; Dana Farber Cancer Institute, Boston, MA, USA; Harvard Medical School, Boston MA, USA; Department of Obstetrics and Gynecology, University of Turku and Turku University Hospital, Turku, Finland

## Abstract

L1 retrotransposons are the only protein-coding active transposable elements in the human genome. Although silenced during normal conditions, they are highly expressed in human epithelial cancers including high-grade serous ovarian cancer (HGSC), where they transcribe to form L1 mRNA and subsequently integrate into the genome by a process called retrotransposition. Despite of high L1 protein expression in the earliest phases of HGSC, these tumors do not accrue many somatic L1 insertions. To understand this unexplained disconnect, we monitored the transcription and retrotransposition activity of two frequently expressed retrotransposition-competent (RC)-L1 (RC-L1) in 64 clinical tumor specimens from 34 HGSC patients and found that despite the presence of RC-L1 mRNA, a third of samples did not acquire somatic L1 insertions. In addition to high inter-patient variability in retrotransposition frequency, there was remarkable intra-patient heterogeneity in L1 insertion patterns between tumor sites, indicating that L1 retrotransposition is highly dynamic *in vivo*. Comparison of genomic and transcriptomic features of L1-null tumors with L1-high tumors (those with ≥5 somatic L1 insertions) showed that retrotransposition was favored by increased rate of cell proliferation.

## Introduction

<u>L</u>ong <u>IN</u>terspersed <u>E</u>lement-1 (LINE-1 or L1) retrotransposons are the only active, protein-coding family of transposable elements in the human genome. They move via a copy-and-paste mechanism called retrotransposition, wherein a 6-kb long full-length retrotransposition competent (RC) L1 transcribes to polyadenylated L1 mRNA which in turn is inserted into the genome by reverse transcriptase activity of L1 proteins encoded by the L1 transcript. As a result of this self-copying activity of L1 retrotransposons, approximately 500,000 copies of L1 sequences are scattered around the genome making up 17% of the human genome (Lander et al., 2001). However, not all L1 copies are retrotransposition-competent (RC) due to inactivating truncation and point mutation accrued during evolution; there are just over 100 RC-L1s in the reference human genome (Beck et al., 2010; Brouha et al., 2003). In fact, evolutionarily young L1s, also called “hot” L1s contribute to the majority of *de novo* L1 retrotransposition events(Beck et al., 2010; Brouha et al., 2003; Rodriguez-Martin et al., 2020; Tubio et al., 2014).

Generally silenced in normal adult cells, L1s are highly active in different epithelial cancers as assessed by the expression of L1 protein L1 ORF1p (Ardeljan et al., 2017; Rodić et al., 2014) and tumor-specific somatic L1 insertions in the cancer genome (Lee et al., 2012; Nguyen et al., 2018; Pitkänen et al., 2014; Rodriguez-Martin et al., 2020; Schauer et al., 2018; Shukla et al., 2013; Tubio et al., 2014). Assessment of L1 activity at the transcriptomic level using bulk RNA- sequencing data in cancer, however, is limited due to technical challenges of reliably detecting L1 transcripts (Lanciano and Cristofari, 2020). This is primarily because L1 sequences are ubiquitously present in the human genome and thus transcripts that contain L1 sequences may not always report on transcription of a full-length L1. For example, if an L1 sequence is present in the intron of an expressed gene, it can be detected as unspliced pre-mRNA and thus can contribute to RNA-sequencing reads (Deininger et al., 2017; Lanciano and Cristofari, 2020). Moreover, the high sequence similarity of young L1 sequences makes it challenging to ascertain which, if any, hot L1 loci with high retrotransposition potential are transcriptionally active in a particular tumor sample.

These challenges have been addressed by innovative bioinformatics methods developed recently (Deininger et al., 2017; McKerrow and Fenyö, 2020; Philippe et al., 2016). However, the transcriptional activity of a RC-L1 does not directly associate with somatic L1 retrotransposition, that is, *de novo* L1 insertions originating from the same source RC-L1, in human cancers. This disconnect is more pronounced in high-grade serous ovarian cancer (HGSC) where L1 ORF1p is highly expressed from very early stages (Pisanic et al., 2019; Rodić et al., 2014; Xia et al., 2017) and yet displays modest somatic L1 retrotransposition events (Rodriguez-Martin et al., 2020). Molecular mechanisms that explains restriction of L1 retrotransposition in HGSC tumors despite high expression of L1 precursors is missing.

In this study, we utilized methods (Deininger et al., 2017; Philippe et al., 2016) that take advantage of 3’
s transduction (**Figure 1A**), a phenomenon by which L1 transcribes and subsequently inserts a non-repetitive sequence on its 3′ flank, to detect L1 transcripts from high-throughput sequencing data. This allowed the detection of frequently expressed RC-L1 loci in high-grade serous ovarian cancer (HGSC) (**Figure 1B**). We screened more than 60 clinical tumor specimens from three different anatomical locations – omentum, ovary and/or adnexal mass of 34 HGSC patients for transcriptional activity of two active source RC-L1 loci. We simultaneously cataloged somatic L1 insertions stemming from these source RC-L1 in the genomes of the same tumor samples using LDI-PCR followed by Nanopore sequencing (Pradhan et al., 2017; Pradhan and Kauppi, 2019) (referred to as LDI-PCR/Nanopore-seq). We found that the presence of an RC- L1 mRNA precursor did not result in high efficiency retrotransposition. On the other hand, we found somatic L1 insertions from an RC-L1 locus in tumors lacking the corresponding RC-L1 mRNA, indicating that transcriptional activity of RC-L1s is dynamic during tumor development. Furthermore, we utilized genomic and transcriptomic data obtained from these HGSC tumor samples and discovered that L1 retrotransposition were more frequent in highly proliferating tumor cells.

**Figure 1.**
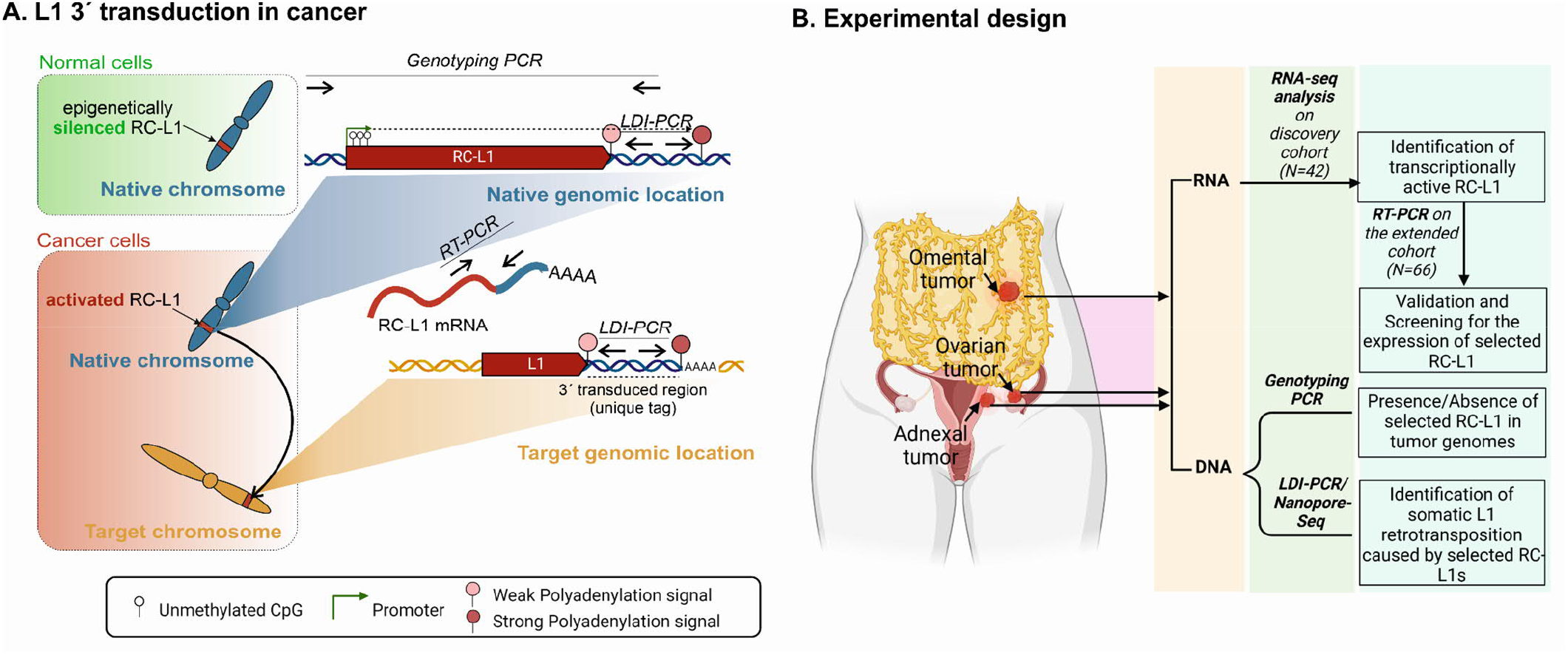
Overview of the study. A) L1 3’s transduction process. Transcriptionally active RC-L1 (present in blue chromosome) generates L1-mRNA including the 3’
s flanking region as they skip past their canonical PAS (pink balloon) and terminates in stronger PAS (red balloon). Such transcript upon successful retrotransposition (in yellow chromosome) integrates non-repetitive flanking region in addition to often truncated L1 sequence. Location of PCR primers (illustrated as arrows showing its 5′ to 3′ orientation) for RC-L1 Genotyping PCR, LDI-PCR and RT-PCR are shown. **B) Study design**. RNA and DNA were simultaneously extracted from the tumors sampled from different anatomical sites from a high-grade serous ovarian carcinoma (HGSC) patient and were subsequently used to detect and screen for transcriptionally active retrotransposition competent (RC)-L1s and identify somatic L1 retrotransposition events that they cause by LDI-PCR and Nanopore sequencing. Genotyping PCR was done to assess the presence or absence of the source RC-L1s in the genomes of the patients.

## Results

### Transcriptionally active retrotransposition-competent L1s in HGSC

Although there are more than 100 retrotransposition competent (RC)-L1s, not all are present in an individual, and furthermore, not all RC-L1s are expressed in a particular cell type or cancer (Philippe et al., 2016). To identify which RC-L1s were active in real-world clinical HGSC specimens, we examined RNA-sequencing data in a discovery cohort of 40 tumor samples from 26 patients with advanced HGSC (**Table 1**) and 4 independent normal fallopian tubes (NFT) samples for expression of 140 intact RC-L1s from L1Base (Penzkofer et al., 2017, 2005) that were also present in the full-length L1 lists of Deininger et al. (2017) (Deininger et al., 2017). Only those L1 transcripts containing a 3’
s flanking region due to transcription read-through, referred to as a 3’ transduced region (**Figure 1A**), are detected by our approach (see Methods and **Figure 2A**).

**Table 1:** Clinicopathological features of the study cohort

|  | **Discovery cohort (N=26)** |  | **Extended cohort (N=34)** |  |
| --- | --- | --- | --- | --- |
| Age at diagnosis | Median (range) | 68 (41–81) | Median (range) | 68 (41–82) |
|  | ≤68 | 13 | ≤68 | 17 |
|  | ≥69 | 13 | ≥69 | 17 |
| FIGO stage | IIIA1 | 1 | IIIA1 | 1 |
|  | IIIB | 0 | IIIB | 1 |
|  | IIIC | 17 | IIIC | 23 |
|  | IVA | 4 | IVA | 4 |
|  | IVB | 4 | IVB | 5 |
| Surgery strategy | NACT | 13 | NACT | 14 |
|  | PDS | 13 | PDS | 20 |
| Residual tumor after surgery | R0 | 13 | R0 | 17 |
|  | non-R0 | 11 | non-R0 | 15 |
|  | n.a. | 2 | n.a. | 2 |
| Response to primary therapy | Complete response | 15 | Complete response | 19 |
|  | Partial response | 7 | Partial response | 8 |
|  | Stable/Progressive disease | 4 | Stable/Progressive disease | 6 |
|  | No Chemotherapy | 0 | No Chemotherapy | 1 |

Abbreviations & Definition: FIGO, Federation of Gynecology and Obstetrics; NACT, neoadjuvant chemotherapy; PDS, primary debulking surgery; R0, surgical resection where no gross or microscopic tumors were left behind.

**Table 2:**
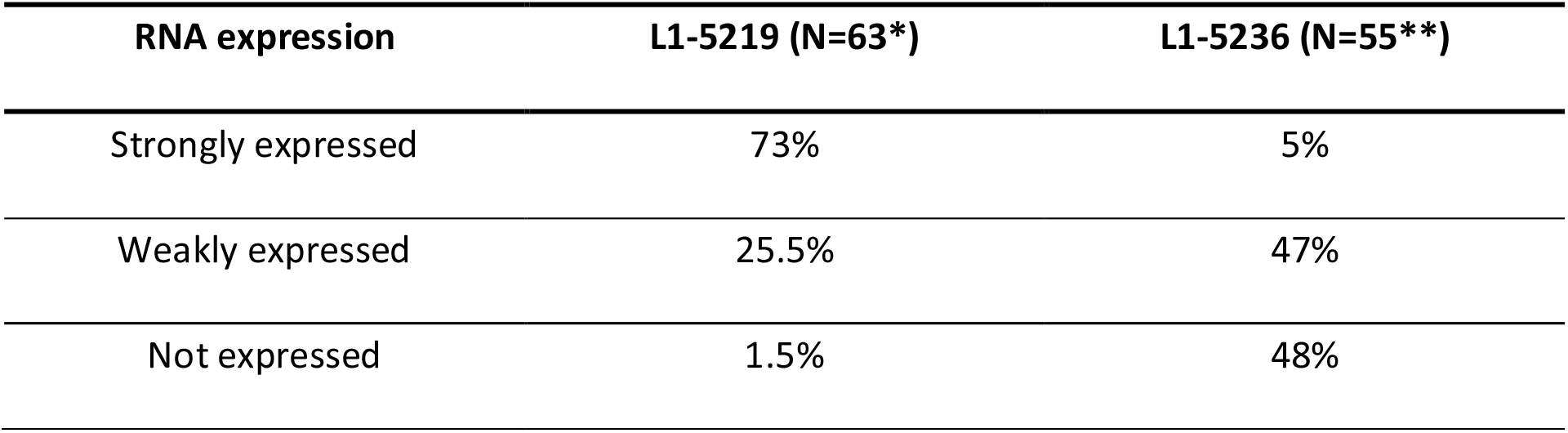
RNA expression of two hot L1s in clinical HGSC specimens, as determined by RT-PCR. (*=N does not include 1 sample where RNA deteriorated; **=N does not include 3 samples where gDNA contamination obscured results and 6 samples that did not have L1-5236 in their genome)

**Figure 2:**
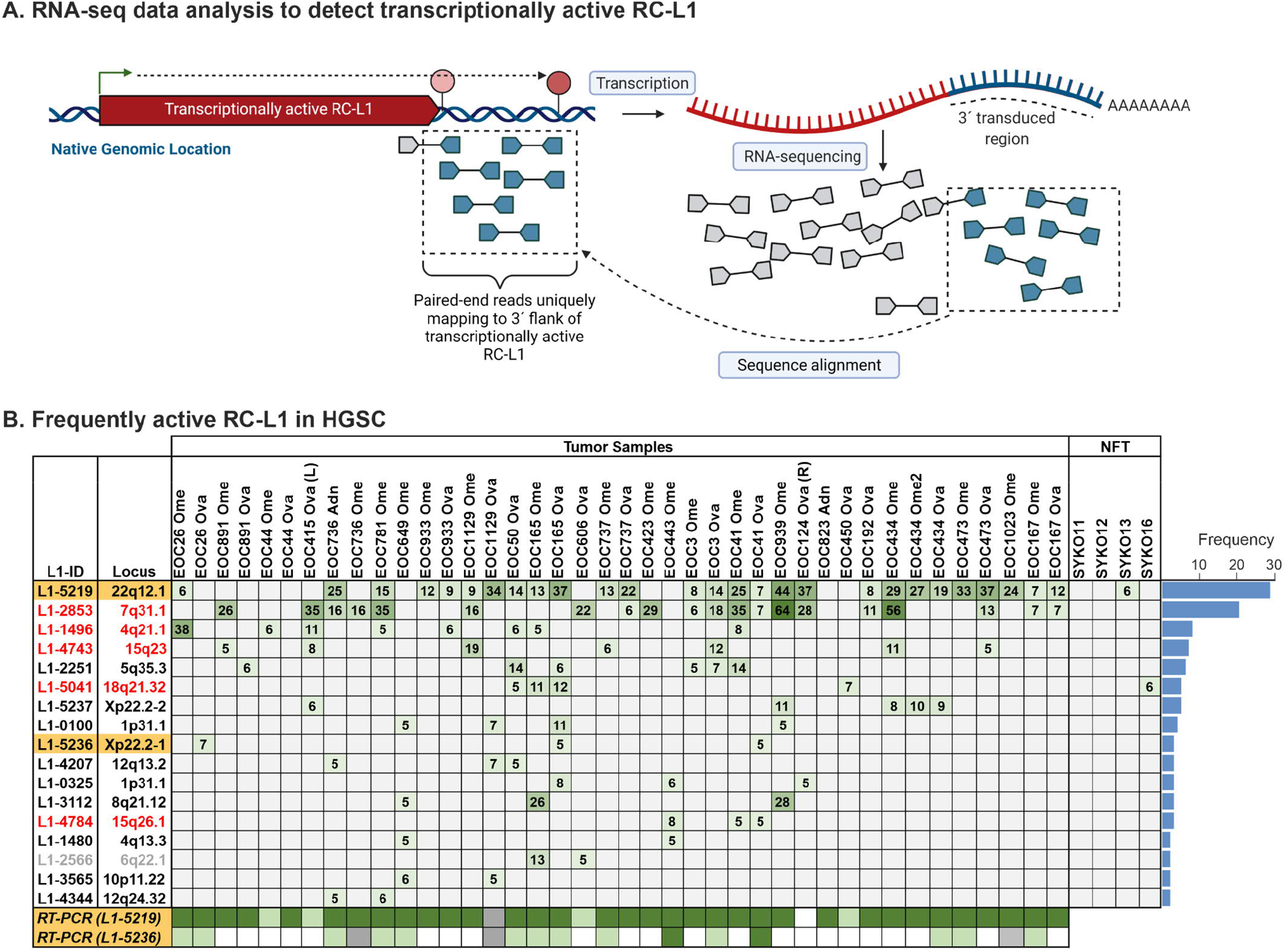
Determination of transcriptionally active RC-L1s. A) Overview of transcriptomic analysis used to detect the presence of RC-L1 transcripts with a 3’s transduced region. Illustration shows short reads (150-300 bp) generated from a RNA-sequencing experiments which are mapped to the reference genome. Reads originating from RC-L1 repetitive sequence (in gray) cannot be accurately mapped, but those that originate from the non-repetitive 3’
s flanking region (in blue) can be mapped to detect a transcriptionally active RC-L1 locus. **B) Full-length retrotransposition competent (RC) L1s expressed in more than one tumor sample**. *In silico* analysis of RNA-sequencing data revealed several RC-L1s (L1-ID as assigned by Deininger et al., 2017) that were expressed in more than 1 HGSC tumor sample. Chromosome locus of each RC-L1 identified is mentioned. The number of transcript fragments detected on the 3′ flank of each RC-L1 in the tumor and normal fallopian tube (NFT) samples is provided in cells. Blank cells indicate that either a) no transcripts were found on the 3′ flank of the RC-L1 in the given sample or b) that the RC-L1 expression did not meet the required cutoff (more in the Methods section). The ID of two selected L1s for validation and LDI-PCR design are highlighted in yellow. RT-PCR validation results for the two selected RCL1s (L1-5219 and L1-5236) are represented by different colors (Dark green= strongly expressed, Light green= weakly expressed, White= Not expressed, and Light gray = not determined due to genomic DNA contamination). RC-L1s that are located inside or nearby a gene in the same orientation as the gene are labeled in red font. Expression of L1-2566 (in gray font) is obscured by presence of another full-length L1 in its vicinity.

L1-5219, a full-length RC-L1 present in the 22q12.1 locus was expressed in 27/40 (68% of) HGSC samples and 1/4 of normal Fallopian tube samples (**Figure 2B**). This source RC-L1 present in the first intron of *TTC28* gene is of particular interest as it is highly mobile in different types of cancers (Cajuso et al., 2019; Nguyen et al., 2018; Pitkänen et al., 2014; Pradhan et al., 2017; Rodriguez-Martin et al., 2020; Tubio et al., 2014), including HGSC. There were several other RC-L1s expressed in more than two HGSC samples, of which we selected L1-5236, located at chromosome Xp22.2, in addition to L1-5219 for downstream assay development. L1-5236 was expressed in 3/40 (8% of) HGSC samples (**Figure 2B**) and was previously reported as actively mobile in HGSC (Nguyen et al., 2018; Rodriguez-Martin et al., 2020). In fact, L1-5236 was identified as one of the highest contributors to 3’
s transduction in ovarian cancer (Rodriguez-Martin et al., 2020). Many frequently expressed RC-L1s identified in our analysis were present within introns that were in the same orientation as the RC-L1s (**Figure 2B**), leading to ambiguity about their L1-promoter specific expression and thus were not considered for downstream assay design. Others were excluded because it was not possible to design specific PCR primers on the flanking 3′-transduced region.

RC-L1s are known to be highly polymorphic in the human population (Beck et al., 2010), thus it is important to first ascertain whether these source L1s exist in the tumor genomes assayed. We examined the presence/absence of the two selected RC-L1s (L1-5219 and L1-5236) in the genomes of an extended HGSC sample cohort, which, in addition to the 40 tumor samples in the RNA-sequencing discovery cohort, included another 24 to make a total of 64 tumor samples from 34 HGSC patients. Genotyping PCR assays (**Figure 3A**) specifically designed for the selected RC-L1s revealed that the allele frequencies of L1-5219 and L1-5236 in our HGSC patient cohort were 1 and 0.75 respectively (**Figure 4A, Supplementary Figure 1**). While three patients’ samples (EOC44, EOC450 and EOC192) lacked L1-5236 entirely, another 12 patients had only one copy of this RC-L1 in their genomes.

**Figure 3.**
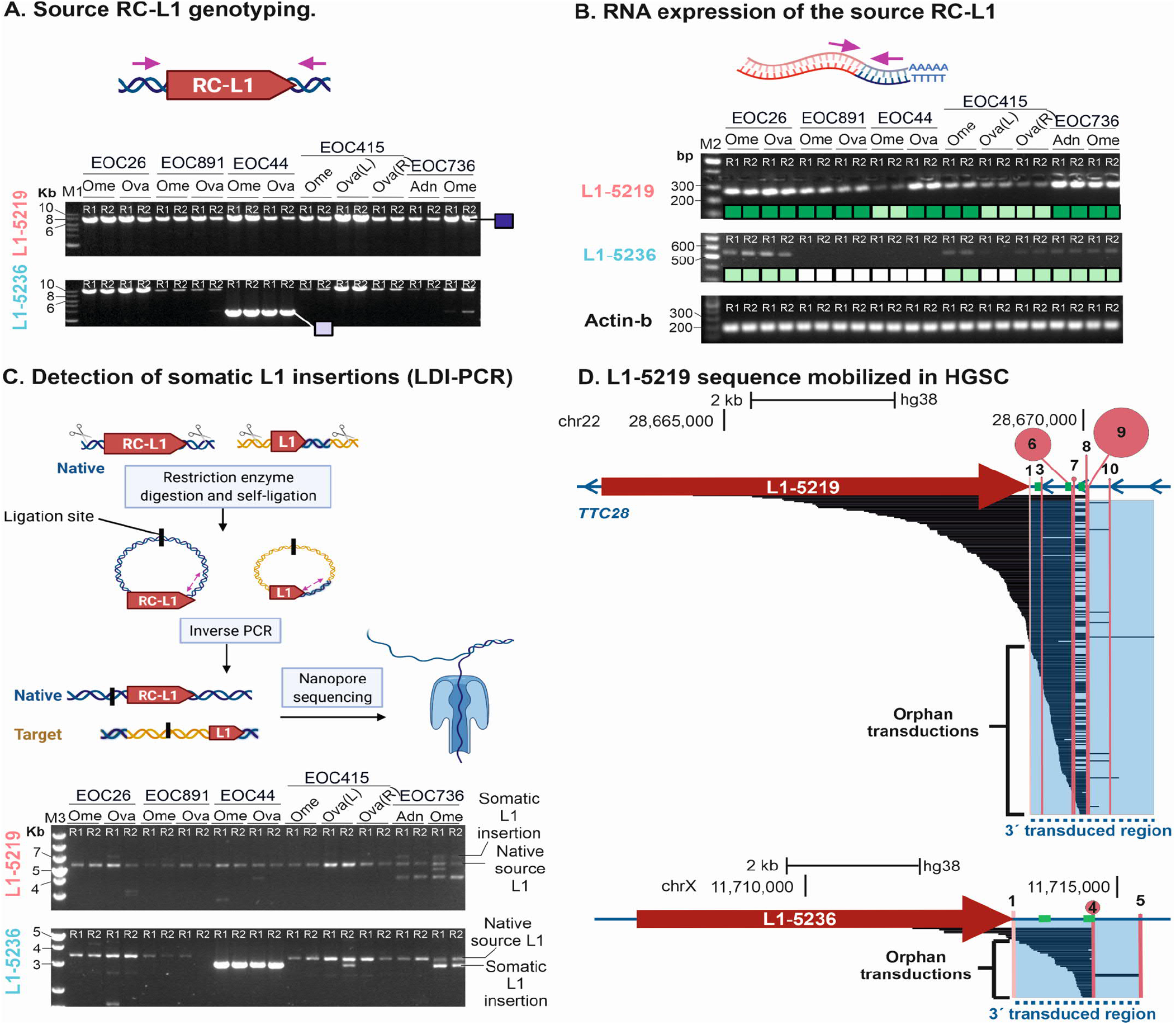
Schematics and representative results of different PCR and sequencing assays to capture different stages of L1 retrotransposition. **A) Genotyping PCR** to determine the presence or absence of the source RC-L1 in the genome. Representative agarose gel images of the genotyping PCR for both RC-L1s, L1-5219 and L1-5236, are shown. Different PCR product sizes show if RC-L1 is present (larger PCR product labeled by a dark blue square) or absent (smaller PCR product labeled by a pale purple square) in the tumor genome. Both PCR products are observed when RC-L1 is present in one copy (heterozygously present; labeled by lilac squares in Figure 4A) **B) Reverse Transcriptase (RT)- PCR** to assess the transcriptional activity of RC-L1. A representative agarose gel image for RT-PCR specific to L1-5219 and L1-5236 locus is shown. Qualitative scoring of RC-L1 expression was done and labeled by dark-green, light-green and white boxes representing strong, weak, and no expression respectively. Expression of the *ACTB* gene was used as a reference for RNA integrity. **C) Long-Distance Inverse (LDI)- PCR/Nanopore sequencing** to identify somatic L1 retrotransposition. Representative agarose gel image of LDI-PCR results for both L1-5219 and L1-5236 locus shows a consistent PCR product in all samples originating from the native RC-L1 and PCR products of different sizes that are unique to a particular patient/tumor sample originating from new somatic copies of RC-L1. These PCR products are subsequently sequenced by Nanopore sequencing to identify the genomic location of somatic copies of RC-L1. (R1 and R2 are two replicates of the same reaction. M1 = 1kb DNA ladder (New England Biolabs); M2 = 100bp DNA ladder (ThermoScientific); M3 = 1 kb plus DNA ladder (ThermoScientific)) **D)** Nucleotide sequences originating from L1-5219 and L1-5236 that are somatically inserted into different genomic locations are plotted below their native genomic location as horizontal black lines. The transparent blue box highlights the non-L1 sequence on the 3’s flank of respective RC-L1s that are mobilized with or without (orphan transductions) L1 sequence. Pink vertical lines show the polyadenylation sites (PAS) downstream of respective RC-L1s that were used for transcriptional termination in the somatic L1-insertions identified in this study with the numbers indicating their respective order in 5′ to 3′ direction with no.1 being RC-L1’s canonical PAS (light pink vertical line). The frequency of PAS usage is depicted by the size of pink circles. Locations of inverse primer pairs are indicated by green lines.

**Figure 4:**
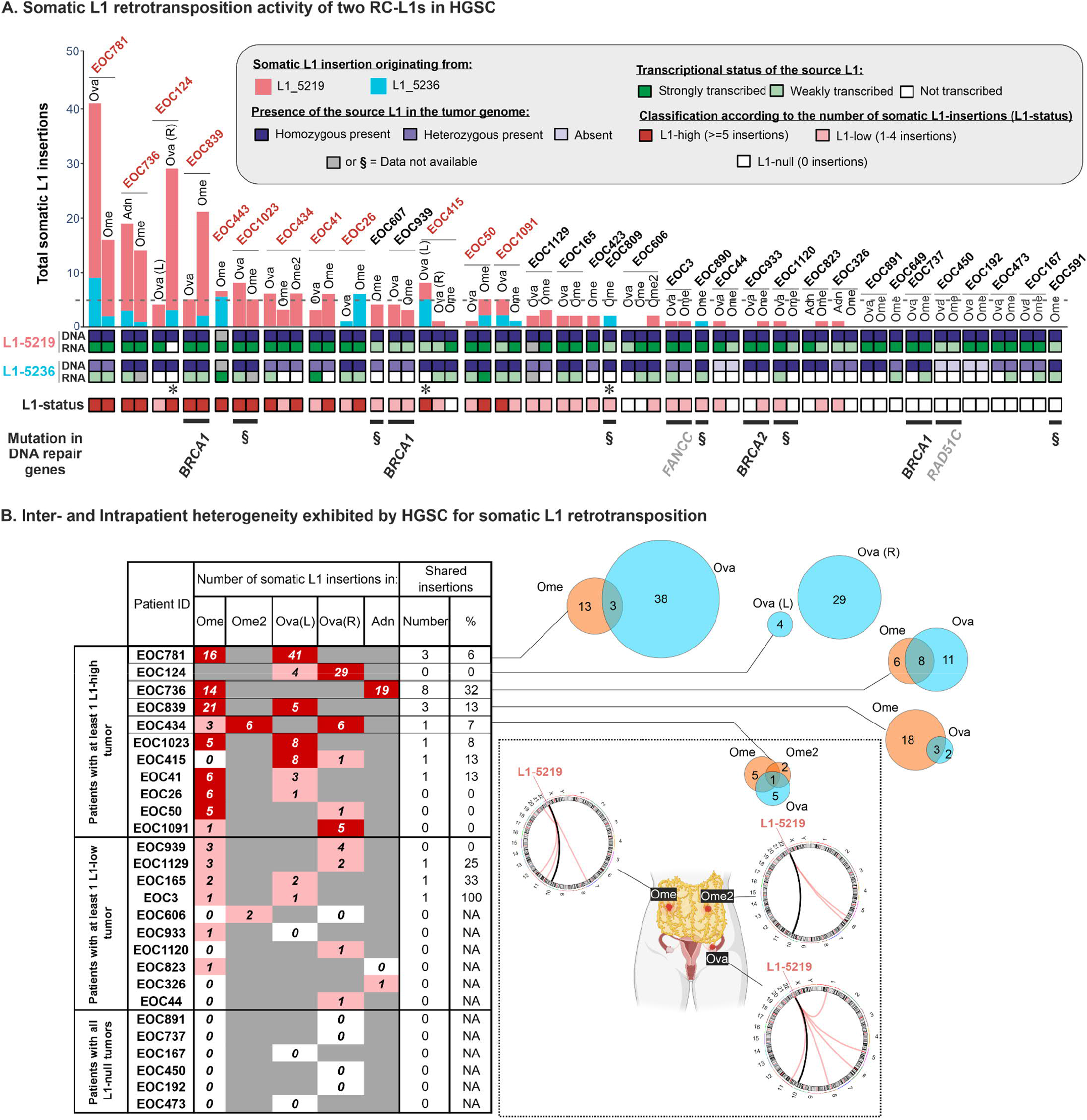
A) Frequency of somatic L1 insertions stemming from two frequently expressed RC-L1s - L1-5219 and L1-5236 in HGSC tumors. The total number of somatic L1 insertions acquired from L1-5219 (red bars) and L1-5236 (blue bars) in tumor samples from the labeled HGSC patients are plotted in a stacked barplot. HGSC patients are in a descending order based on average somatic L1 insertions found in their tumors and patients with at least one L1-high tumor (tumors with ≥5 L1-insertions) are labeled in red font. Anatomical location of the tumors are labeled as “Ome” for omentum, “Ova” for ovary, and “Adn” for adnexa (In cases when ovarian tumors were sampled from both ovaries of a patient, it was specified if they were found on the left (L) or right (R) ovary. When more than one omental tumor were sampled from a patient, serial numbers were assigned to each tumor). The squares below the stacked barplot show the genomic (DNA) and transcriptomic (RNA) status of the source L1 investigated. Gray boxes indicate that either the PCR was unsuccessful or in the case of RT-PCR, the results were compromised due to likely genomic DNA contamination. Asterisks (*) indicate tumor samples in which somatic L1 insertions are identified despite the corresponding source L1 being transcriptionally inactive. HGSC tumors were classified as L1-high (≥5 somatic L1 insertions), L1-low (1-4 L1 insertions) and L1-null (no L1 insertion) and labeled with red, pink and white boxes respectively. Somatic (black font) and Germline (gray font) mutations observed in HGSC tumors and patients are indicated in a row below (§, WGS data not available). **B) HGSC tumors display high levels of intra-patient heterogeneity in terms of somatic L1-insertions**. The total number of somatic L1 insertions in different tumor samples from HGSOC patients are listed along with the total number and percentage of shared insertions between tumors from different sites. Intrapatient heterogeneity is also illustrated by Venn diagrams that represent different tumor sites for 5 patients with most somatic L1 insertions. In the inset variability of somatic L1-insertion profile in two tumors located at omentum and one on the ovary of patient EOC434 is demonstrated using Circos plots that join the native RC-L1 with the genomic location of their somatic L1 copies. Red links show unique somatic L1-insertions in each tumor originating from L1-5219 (chr22). Black links show one shared somatic L1-insertion in *ATRNL1* gene common to all the tumors.

To validate RNA-Seq findings, we assayed transcriptional activity of these two RC-L1 loci by RT-PCR in the extended HGSC cohort. RT-PCR results were semi-quantitatively scored as “strongly expressed”, “weakly expressed” or “not expressed” based on results from agarose gel electrophoresis (**Figure 3B, Supplementary Figure 2**). While 98.5% of samples in the extended cohort showed some level of expression of L1-5219, 52% of the tumor samples having at least one copy of L1-5236 in their genomes were positive for L1-5236 expression (**Figure 4A, Table 1**). As RT-PCR was also performed on the same 40 samples whose RNA-Seq data were analyzed to identify transcriptionally active RC-L1, we were able to compare the sensitivity of detection between the two approaches (**Figure 2B**). RT-PCR proved to be a more sensitive approach overall. In addition to detecting transcriptional activity of L1-5219 and L1-5236, thereby confirming RNA-Seq results in all but one case (no activity of L1-5219 in EOC124 Ova(R)), RT-PCR also detected their expression in samples that were negative for L1-5219 and/or L1-5236 expression by RNA-Seq *in silico* analysis (**Figure 2B**).

In summary, we identified transcriptionally active RC-L1s in HGSC using RNA-Seq data obtained from 40 tumor samples (discovery cohort) and selected two RC-L1s, L1-5219 and L1-5236, previously shown to actively mobilize in HGSC, for downstream assays. We then screened for the genomic presence and transcriptional activity of these two RC-L1s in a larger sample cohort. While L1-5236 was more polymorphic (allele frequency = 0.75) and less expressed in our sample cohort, L1-5219 was almost fixed and expressed in nearly all tumor samples.

### Somatic L1 insertions caused by two frequently expressed RC-L1s in clinical HGSC samples

To establish whether the transcriptionally active loci L1-5219 and L1-5236 give rise to somatic L1 insertions in HGSC genomes, we assayed tumor DNA of the same samples for L1 retrotransposition by LDI-PCR/Nanopore-seq. We detected 233 unique *de novo* L1 insertions arising from these two RC-L1 loci, out of which 190 were contributed by L1-5219 and 43 by L1- 5236 (**Figure 4A** and **Supplementary Table 1**). 42/64 samples (66%) showed at least one somatic L1 insertion, and 17 had ≥ 5 insertions (classified as L1-high samples, **Figure 4A**). To ensure that the L1 insertions detected in the tumor samples were somatic and not germline events, we performed LDI-PCR on the matching normal genomic DNA obtained from the patients’ blood samples. While tumor samples showed reproducible LDI-PCR products of sizes distinct from the “native” RC-L1 specific product indicating *de novo* L1-insertions, the matching normal samples only produced native RC-L1 specific product (**Supplementary Figure 3**) hence confirming the somatic nature of the L1 insertions detected in tumor DNA.

22 samples did not acquire any somatic L1 insertions and were hence classified as L1-null samples. All of these samples were positive for transcriptional activity of at least one of the two RC-L1 loci (**Figure 4A;** EOC124 Ova(R) was an exception where expression of neither RC-L1 was detected by RT-PCR despite showing expression of L1-5219 in RNA-Seq analysis, see **Figure 2B**). This finding demonstrates that the presence of RC-L1 mRNA does not guarantee their insertion into the genome.

All 6 samples from 3 patients who lacked L1-5236 altogether did not generate any somatic L1 insertions from this locus, as expected. However, in three samples where L1-5219 and/or L1- 5236 transcripts were undetectable by RT-PCR, somatic L1 insertions stemming from these loci were present (**Figure 4A**). This implies that L1-5236 was active during tumor evolution and the detected insertions represent historical events that occurred before the locus was (again) transcriptionally silenced.

We then asked if the number of somatic L1 insertions in tumors from the same patient differed based on their anatomical location. There were 19 cases where we had sampled tumors from two distinct anatomical regions, one from Omentum and another from Fallopian Tube-Ovary (FTO). HGSC typically originates in the fallopian tube and primary tumors develop in the ovaries, while tumors in the omentum represent metastases. Pairwise comparison for each case did not show any proclivity towards tumors found in a particular anatomical site (**Supplementary Figure 4**).

Long-read sequences generated from LDI-PCR/Nanopore seq capture the sequence of the source or parent RC-L1 as well as their novel offspring copies in their entirety. This allowed detailed characterization of the somatic L1 copies (explained in detail in **Supplementary Result 1**) revealing *i)* a high incidence of orphan transduction events in HGSC where the offspring copy is truncated to the extent that the insertion contains no L1 sequence at all (**Figure 3D**) and *ii*) the most preferred polyadenylation signal (PAS) downstream both RC-L1s for L1 transcriptional termination (**Figure 3D**).

Although 116 insertions (50%) occurred in introns of protein-coding or long non-coding RNA genes, the majority were present in genes that were expressed at very low levels in HGSC tumors (**Supplementary Figure 5**), in line with previous studies (Jung et al., 2018; Tubio et al., 2014). In two genes that were substantially expressed in HGSC tumors, somatic L1 insertion seemed to have two opposite effects (explained in detail in **Supplementary Result 2**). Both up- and as down- regulating effects of somatic L1 insertion in the host gene have been reported (Nguyen et al., 2018; Shukla et al., 2013). It should be noted that our results may be obscured by host gene expression originating from other cell types, e.g. fibroblasts, and/or from other tumor cells that are negative for that particular insertion. The latter is an important consideration as our results indicate that somatic L1 insertions detected by our assay are subclonal in nature (explained in detail in **Supplementary Result 3; Supplementary Figure 6 & 8**).

Altogether, we were able to identify 233 insertions in 64 HGSC samples using a highly sensitive approach to detect somatic L1 insertion stemming from two frequently expressed RC-L1s. Integrating data that report on the transcriptional status of RC-L1 as well as retrotransposition from the same tumor sample, we were able to infer that L1 transcriptional status can switch during tumor evolution. We also show that transcriptional activity of RC-L1 does not guarantee somatic L1 retrotransposition. Furthermore, anatomical location of tumors had no significant impact on the number of somatic L1 insertions acquired.

### High inter- and intra-patient heterogeneity in somatic L1 retrotransposition profiles

We observed high inter-patient heterogeneity with regards to total unique somatic L1 retrotransposition events. The range was 0-54 insertions; based on the number of somatic insertions, we classified the L1 insertion status of each sample as either L1-null (0 insertions), L1- low (1-4 insertions) or L1-high (5 or more insertions). Although we have profiled retrotransposition arising from two highly active RC-L1, it should be noted that these L1-null samples may contain insertions stemming from other RC-L1 and/or retrotransposition events without 3’ transductions that are not detected by our assay. 22 out of 64 tumor samples were L1-null (**Figure 4A**). At the patient level, 8 out of 34 patients were L1-null (**Figure 4A**) while 12 patients had at least one L1-high tumor sample. L1 insertion status of different anatomical regions of the same HGSC patient was consistent, with only a few exceptions (**Figure 4A**). L1- insertion burden in the tumor of the patients were not associated with their clinical features such as age, primary therapy outcome or tumor stage (details in **Supplementary Result 3 and Supplementary Figure 8**)

For 27 patients assayed for somatic L1 retrotransposition, we had tumor samples from at least two anatomical sites. While the number of somatic L1 insertions found in different tumors from the same patient or their L1 status (**Figure 4**) was comparable, their insertion profile was highly variable (**Figure 4B**). That is, within the same patient, most somatic L1 insertions were unique to each tumor and only a few were shared between the tumors. The percentage of shared insertions between tumor samples from patients with more than 5 total independent events ranged from 0 to 32% (**Figure 4B**). To assess whether this intrapatient heterogeneity may be a result of technical artifacts during LDI-PCR/Nanopore-seq, we validated insertions that were shared between tumor samples and those that were found to be unique to one tumor sample of patient EOC415 using two PCR-based assays (**Supplementary Figure 9**). Somatic L1 insertion-specific PCR products were observed by at least one of the validation PCR assays only in those samples where such insertions were detected initially by LDI-PCR/Nanopore-seq (**Supplementary Figure 9**), indicating that insertional profiles between anatomical sites are distinct.

### L1 insertion burden in HGSC tumors were positively associated with with increased genomic breakpoints and a mitotic clocklike mutational signature

Whole-genome sequencing (WGS) data of 52 tumor samples from 29 patients (**Supplementary Table 2, *cohort description***) and their matching blood DNA were analyzed as a part of a separate study (https://www.deciderproject.eu/). As different host DNA repair factors are predicted to play a role in the L1 retrotransposition process (Goodier, 2016) we checked for germline or somatic biallelic loss-of-function mutations in DNA repair genes. 3 HGSC patients in our cohort with biallelic *BRCA1* loss exhibited three different L1 insertion phenotypes – L1-high, L1-low, and L1-null (**Figure 4A**). Patient EOC933 who carried a somatic *BRCA2* mutation and LOH of the functional allele had one somatic L1 insertion in one of the two tumors sampled. There were two patients with germline mutations in one allele of *FANCC* (EOC3) and *RAD51C* (EOC450) and the functional copy inactivated via LOH during tumor evolution. While one common somatic L1 insertion was found in both tumors from patient EOC3, patient EOC450 displayed no somatic L1 insertions (**Figure 4A**). However, it is not possible to determine which event, somatic L1 insertion or the somatic mutations and/or subsequent LOH occurred first, limiting our interpretation of the role of these DNA repair genes in the L1 retrotransposition process.

We associated the genomic features obtained from WGS analysis with somatic L1 insertion burdens. As samples from the same patient had similar genomic features, we picked one representative sample with the highest number of L1 retrotransposition events per patient to avoid representation biases. We also excluded tumor samples from patients EOC44, EOC450 and EOC192 because they lacked L1-5236 in their genome which may lead to underestimates of their L1 insertional burden. After applying these selection criteria, we used genomic data from 24 tumor samples from 24 HGSC patients to explore their association with L1 insertion burden. As recent studies have shown elevated levels of somatic L1 retrotransposition in aneuploid tumor cells (Rodriguez-Martin et al., 2020; Yamaguchi et al., 2020), we compared the ploidy of tumors that had different L1 insertion burdens (**Figure 5A**). We did not observe significant differences in ploidy when we compared three groups of tumors based on their L1-status (**Figure 5A**). We also compared the total number of genomic breakpoints between samples with different L1-status (**Figure 5B**) and found that L1-high tumors had significantly more breakpoints than L1-low tumors (*p*=0.02, student t-test).

**Figure 5:**
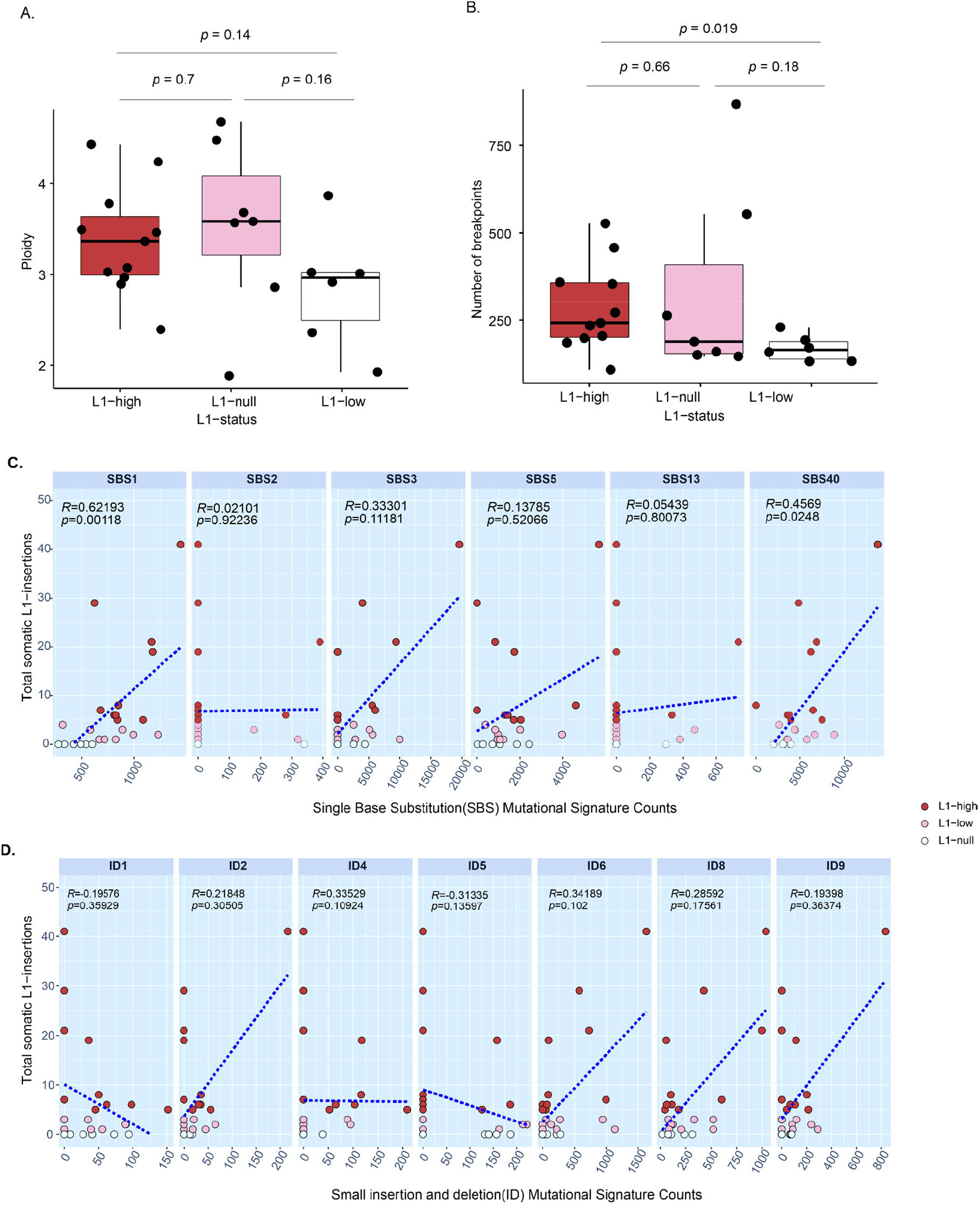
Association of the L1-status of HGSC tumors with ploidy, number of breakpoints, and mutational signatures of their genomes. Ploidy and the total number of genomic breakpoints observed by whole-genome sequencing in L1-high (N=10), L1-low (N=8), and L1- null (N=6) tumor samples are shown in boxplots A and B respectively (p-value obtained after a two-tailed student’s t-test is indicated). Relationship between the somatic activity of hot RC- L1s with Single-base substitution (SBS) and Small insertion and deletion (ID) mutational signatures observed in HGSC tumors (C and D respectively). The total number of somatic L1- insertions detected in HGSC tumors (N=24) is plotted against different indel (ID) and single base substitution (SBS) mutational signature counts obtained by analyzing WGS data from the matching tumor (R and *p*-value calculated by Spearman’s correlation test)

We also analyzed association of single-base substitution (SBS) and short insertion-deletion (ID) mutational signatures (Alexandrov et al., 2020, 2013) with the total number of somatic L1 insertions (**Figure 5C** and **D** respectively). SBS1 and SBS40 were the only mutational signatures that were significantly associated with somatic L1-insertion burden (Spearman’s Rho (R)=0.62 and 0.45; *p*-value=0.001 and 0.02 respectively, **Figure 5C**). SBS1 and SBS40, both, are associated with the age of the individual and SBS1 specifically correlates with the rate of cell divisions and thus acts as a mitotic clock (Alexandrov et al., 2020; Nik-Zainal et al., 2012).

In summary, we show that increased somatic L1 retrotransposition is observed in tumors with higher number of breakpoints. We also found positive correlation of L1 insertion burden with mutational signatures that are associated with higher rates of cell division.

### Polymorphism in the source L1 could inactivate its retrotransposition activity

Almost all (98.5%) tumor samples expressed L1-5219 RNA (**Table 1, Figure 4A**). In 28 tumor samples (44% of the sample cohort), despite the presence of the precursor L1-5219 mRNA, these elements were unable to generate insertions. As L1 mRNA codes for L1 proteins ORF1p and ORF2p that are critical for a successful L1 retrotransposition event, an inactivating mutation in these coding sequences may render the L1 incapable of retrotransposition. Thus we inspected variation in the source L1-5219 sequence of samples with no somatic insertion arising from L1- 5219 (referred to as L1-5219**-** samples) and compared them with those samples that had at least one L1 retrotransposition event (referred to as L1-5219^**+**^ samples). We were able to do this using sequencing data generated by LDI-PCR/Nanopore-Seq, because the source L1 in its native genomic location is sequenced along with the daughter copies of the source L1 generated by L1 retrotransposition (**Figure 3C**).

With the DNA sequencing data available we were able to assess the L1-5219 sequence and ∼2 kb of its 3’
s flanking genomic region. We found several variants that were unique to L1-5219^**-**^ samples in the 5′ UTR and L1-ORF2p coding region as well as in its 3′ flanking region (**Figure 6**). All variants were present heterozygously and two variants (rs201455670 and variant#1, **Figure 6**) were present in the ORF2p coding sequence of L1-5219. The variant rs201455670 observed in L1-5219^-^ patient EOC737 changes threonine at the 655^th^ position of the L1 ORF2p protein to proline. The mutated amino acid is part of the reverse transcriptase domain of L1 ORF2p necessary for L1 retrotransposition and thus conceivably could render the L1-5219 incompetent for retrotransposition (Moran et al., 1996). In fact, a scanning mutagenesis study investigating the effect of the alteration of each amino acid in L1 ORF2p in the retrotransposition efficiency shows that alteration of this amino acid reduces the retrotransposition efficiency by almost 90% (Adney et al., 2019). In addition to 64 clinical HGSC tumor specimens, we also tested the somatic retrotransposition activity of L1-5219 in 7 commercially available HGSC cell lines (**Supplementary Figure 10**). Strikingly, variant rs201455670 (**Figure 6**) was also present in the Kuramochi cell line homozygously. Another variant in ORF2p domain of L1-5219 (variant#1 in **Figure 6**) that was detected in patient EOC26 was synonymous and thus is not expected to lead to any changes in the ORF2p protein domain.

**Figure 6:**
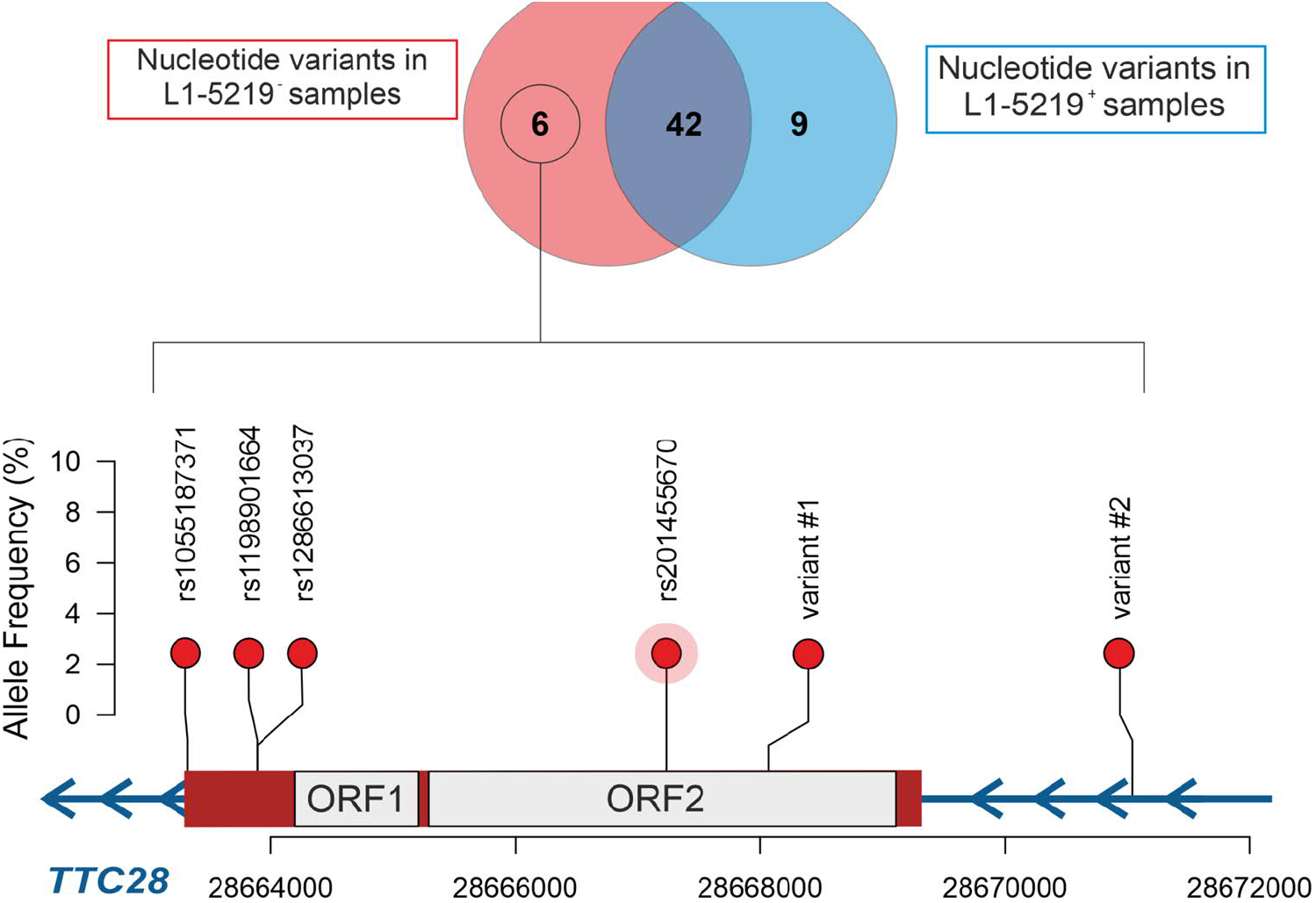
Sequence analysis of source L1-5219 to detect potentially inactivating variants. Out of 58 total SNVs observed in the L1-5219 locus as identified in LDI-PCR/ Nanopore-seq data, 6 were unique to the samples with no somatic L1-insertions originating from L1-5219 (L1-5219^**-**^ group; total=27) and 9 were unique to samples with at least 1 somatic L1-insertion originating from L1-5219 (L1-5219^**+**^ group; total=37). Genomic locations of the variants unique to the L1- 5219^**-**^ group are plotted using red balloons with the height of balloons corresponding to their allele frequency in our patient cohort. ORF1 and ORF2 region of the L1 that encodes L1-ORF1p and L1-ORF2p proteins respectively are annotated in gray boxes. Variants are annotated with their SNP IDs when available, otherwise, they are assigned with a unique serial number. rs201455670, also identified in one HGSC commercial cell line Kuramochi with no somatic L1- 5219 mediated insertion is highlighted.

In summary, mutations in the internal protein-coding region of an RC-L1 may inactivate it and thereby convert it to become retrotransposition incompetent. Mutations in hot RC-L1s thus can obscure the frequency of somatic retrotransposition in tumors of patients that are carriers of such mutations.

### Differential gene expression and Gene set enrichment in L1-high vs L1-low samples

To investigate transcriptomic differences between tumor samples that are either permissive or restrictive to the L1 retrotransposition process, we compared RNA-Seq data from L1-high and L1- null tumor samples, i.e. those that had ≥5 versus no somatic L1 insertions, respectively. In the L1- null group, we only considered those samples that had at least one functional copy of both source L1s. For instance, tumors from patients EOC44, EOC450 and EOC192 were excluded from the analysis despite being L1-null, as L1-5236 was not present in their genome. In total, RNA-Seq data was available for 11 samples in the L1-high and L1-null groups each (**Supplementary Table 2**). All samples had tumor purity of more than 20%, as measured by allele frequency of clonal *TP53* mutation. Furthermore, the bulk transcriptomic data was deconvoluted for cancer-cell specific expression (Häkkinen et al., 2021).

We performed differential gene expression analysis and identified several genes that were significantly upregulated in the L1-high group when compared to the L1-null group (**Figure 7A, Supplementary Table 3**) – RNA expression of *Fibroblast Growth Factor* 3 (*FGF3*) and *Polymeric Immunoglobulin Receptor* (*PIGR*) were the most significantly upregulated gene in L1-high and L1- null group, respectively. Both of these genes, have however not been associated with L1- retrotransposition regulation. Out of all the genes that were significantly upregulated in either L1-null or L1-high tumors, only *LIN28A* (upregulated in L1-high tumors), to our knowledge, can be attributed to L1-regulation. *LIN28A* is a known repressor of miRNA *let7 (Piskounova et al*., *2011)*which in turn has been shown to negatively regulate L1 retrotransposition (Tristán-Ramos et al., 2020).

**Figure 7:**
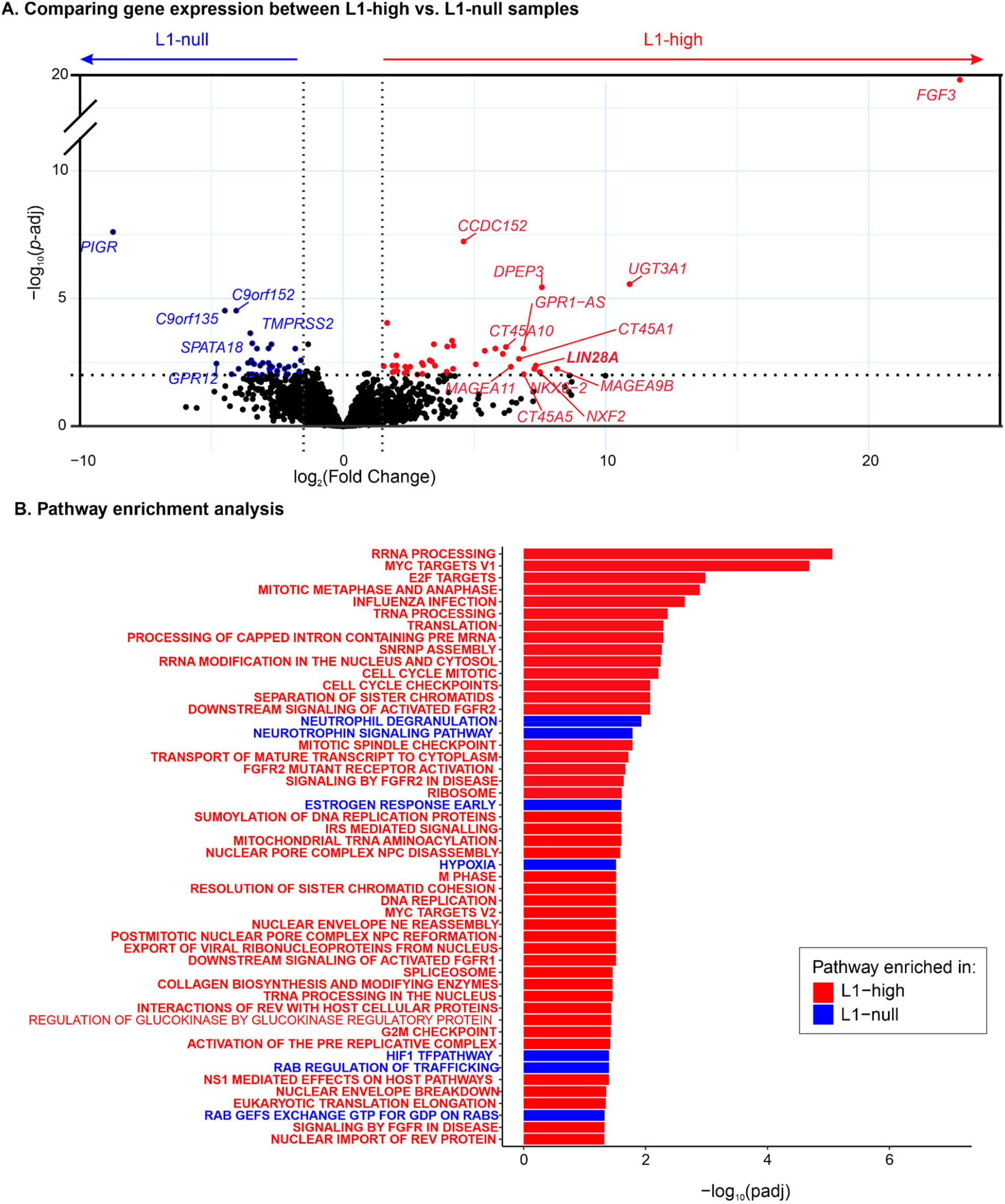
Comparison of the transcriptomic profile of L1-high and L1-null HGSC tumors. A) Significantly upregulated genes in L1-high samples (n=11) and L1-null samples (n=11) as discovered by differential gene expression analysis are represented by red and blue circles respectively in a volcano plot. The cutoff for False Discovery Rate (FDR) or padj<0.01 and Fold change>1.5. Highly upregulated genes in both groups are labeled with gene names. B) Pathways significantly enriched in L1-high samples (red) and L1-null samples (blue) as determined by Gene Set Enrichment Analysis (GSEA) are plotted in a descending order based on their false-discovery rate (FDR). Pathways with FDR<0.05 are shown in the barplot.

To assess which pathways were enriched in each group we performed a Gene-Set Enrichment Analysis (GSEA) of differentially expressed genes between the two groups (**Figure 7B**). Gene-sets enriched in the L1-high group included those involved in pathways related to mitotic progression, DNA replication, and RNA processing (FDR<0.05, **Figure 7B**). Enrichment of pathways involving cell cycle progression in L1-high tumors is in line with our finding that mitotic clock-like mutational signature SBS1 occurred more in tumors with high somatic L1-insertions (**Figure 5C**). Moreover, the age of patients, which also causes accumulation of the aforementioned signatures, did not show any association with the number of somatic L1 insertions acquired (**Supplementary Figure 8A**) indicating that not age *per se* but the rate of tumor cell division that affects the number of somatic L1 retrotransposition events in HGSC.

In the L1-null group, on the other hand, pathways related to hypoxia and response to hypoxia such as the HIF1 pathway, as well as genes involved in neutrophil degranulation, early estrogen response, and neurotrophin signalling were enriched (FDR<0.05, **Figure 7B**). High immune activity has been associated with low somatic L1 retrotransposition activity in gastrointestinal cancers (Jung et al., 2018). In these cancer types, the immune activity was believed to be elicited by Epstein-Barr Virus infection and microsatellite instability. As hypoxia and the subsequent activation of HIF1 pathway are also known to elevate expression of inflammatory genes (Corcoran and O’Neill, 2016), it is possible that in HGSC, hypoxia-induced inflammation leads to suppressed L1 retrotransposition activity.

## Discussion

In this study, we identified RC-L1s that are transcriptionally active in high-grade serous ovarian cancer (HGSC) and chart their expression and somatic L1 retrotransposition profiles in 64 tumor samples from 34 patients. While we observed high variability between HGSC patients in terms of total *de novo* somatic L1 insertions acquired, different tumors from the same patient varied in their somatic L1 insertion profile, that is, there are only a few insertions shared between different intra-patient tumor sites. There are several factors that contributed to the interpatient variability. First, interpatient variability can arise due to the lack of evolutionarily young RC-L1s in the patient’s genome. L1-5236, one of the two RC-L1 selected for our study, was completely absent in 3 out of 34 patients. Second, lack of retrotransposition in some patients may be due to inactivating variants in their RC-L1. Despite the presence of a functional copy of RC-L1 in their genomes and subsequent expression of such RC-L1s on an mRNA level, some tumors showed no insertion at all (L1-null tumors) while others generated as many as 54 *de novo* L1 insertions. To understand this variability, we compared the transcriptomes of such L1-null tumors with that of L1-high tumors (tumors that had acquired more than four somatic retrotransposition events). L1- null tumors were enriched for hypoxia and related pathways that could potentially trigger an immune response. L1-high tumors on the other hand were enriched for gene sets that drove mitotic progression. This finding is supported by results obtained after analyzing WGS data from the HGSC tumor samples where we observed a positive correlation between the total somatic L1 insertion with mutational signatures that accumulate as a result of increased cell division.

In addition to our main findings, we were able to characterize each somatic L1 insertion in greater detail by sequencing LDI-PCR products with Nanopore sequencing. By focusing on 3’
s transduction events we found more than 60% to be orphan transductions, meaning they did not contain the L1 repetitive sequence at all. This is an important observation, given that most of the existing targeted sequencing method targets the internal (L1 repetitive) sequence of young RC-L1 (Nguyen et al., 2018; Shukla et al., 2013; Steranka et al., 2019), and thus would miss these insertions entirely. *In silico* methods used to detect somatic L1 retrotransposition from WGS data (Rodriguez-Martin et al., 2020; Tubio et al., 2014) are able to detect these orphan transductions, but they are substantially less sensitive than targeted sequencing methods such as LDI- PCR/Nanopore seq (Pradhan et al., 2017) and hence are unable to identify subclonal events. We were also able to pinpoint preferential polyadenylation signal (PAS) usage for transcriptional termination downstream of both RC-L1’s canonical PAS. This resource can be used in the future to develop locus-specific assays to monitor somatic L1 retrotransposition rates in diseases.

Recent advances in profiling somatic L1 retrotransposition by targeted sequencing and WGS analysis have shown that retrotransposition is prevalent in many epithelial cancer types, once L1 promoters are derepressed leading to transcription of retrotransposition-competent L1s (Cajuso et al., 2019; Lee et al., 2012; Pitkänen et al., 2014; Scott et al., 2016; Shukla et al., 2013; Solyom et al., 2012; Tubio et al., 2014). It is, however, still unclear if the presence of RC-L1 mRNA in these epithelial cancers ensures its integration into the tumor genome. Advanced HGSC is a fitting model to investigate this question as *TP53*, a negative regulator of L1 expression (Wylie et al., 2016) is disrupted in this type of cancer. Accordingly, we observed transcripts of at least one out of two RC-L1s assayed in almost all HGSC tumors assayed. Availability of the RC-L1 mRNA, however, did not guarantee somatic L1 insertions in HGSC. This is in agreement with the literature, according to which more than 90% of ovarian cancer cases (Rodić et al., 2014) showed the presence of the L1 protein L1-ORF1p, but only 60% had at least one somatic L1 retrotransposition (Rodriguez-Martin et al., 2020). L1-ORF1p expression, however, can originate from any out of thousands of L1, including retrotransposition-incompetent L1s in the human genome. Thus our locus-specific examination of RNA and DNA for transcription and retrotransposition activity of specific RC-L1s, respectively, provide direct evidence that somatic L1 retrotransposition in HGSC is regulated at a post-transcriptional level.

Our finding supports a previous study that integrated genomics and transcriptomics data of ∼200 gastrointestinal (GI) cancers (Jung et al., 2018) and showed lower somatic L1 retrotransposition activity in tumors with active immune pathways. While Epstein-Barr virus co-infection or microsatellite instability (MSI) were believed to be the factors that triggered an immune response in these GI cancers, hypoxia and HIF pathway activation that were enriched in L1-null HGSC tumors could elicit the same response in HGSC, making these tumors restrictive of somatic L1 retrotransposition activity. However, we did not find any positive correlation between total somatic L1 insertion and the age of the HGSC patients as Jung et al. (2018) did in GI cancer. We speculate that this could be because *TP53*, a negative regulator of L1 expression (Wylie et al., 2016) is ablated early in HGSC tumorigenesis and thus age-related L1 derepression (Simon et al., 2019; Van Meter et al., 2014) likely is not a prerequisite for somatic L1 retrotransposition in HGSC.

L1-high tumors were enriched for transcriptomic profiles that were related to cell cycle progression. This, along with the fact that the frequency of somatic L1 insertions correlated positively with mitotic-clock-like mutational signature, suggests that more somatic L1 insertions are accrued in tumors that are highly proliferative. *In vitro* studies have shown that cell division is vital for successful L1 retrotransposition as nuclear envelope breakdown provides the L1 RNP complex with access to the nuclear genome (Mita et al., 2018; Xie et al., 2013). Thus our study provides *in vivo* evidence of a positive association of cell division with somatic L1 retrotransposition frequency.

A limitation of our study is that only two RC-L1s out of hundreds that are present in the genome were assessed for transcriptional and somatic retrotransposition activity. We prioritized specific detection of RC-L1 transcripts and somatic L1 insertions originating from these two loci, L1-5219 and L1-5236, over the number of RC-L1 tested. We consider this a justified approach, given that analysis of tumor genomes from 118 ovarian adenocarcinoma patients has shown that the two RC-L1s selected for this study contribute >25% of somatic 3’
s transduction events (Rodriguez-Martin et al., 2020). Thus, although limited to two, the RC-L1s selected appear to be robust *in vivo* reporters of somatic retrotransposition upon transcription.

In conclusion, we provide evidence of post-transcriptional regulation of L1 retrotransposition in HGSC. Using both genomic and transcriptomic information of L1-high and L1-null tumor samples, we show that L1 retrotransposition are more frequent in rapidly proliferating tumor cells. Our findings also suggests that upregulation of genes such as *LIN28A* facilitates L1-retrotransposition in HGSC while upregulation of hypoxia and related pathways deters it.

## Materials and Methods

### Human tumor specimens

Tumor samples from HGSC patients at the Turku University Hospital, Finland prior to any treatment were used for this study. After sampling of these chemonaive tumors, patients either went through Primary debulking surgery (PDS) or Neoadjuvant chemotherapy (NACT). We focused on samples primarily taken from two anatomical regions - Omentum and Fallopian Tube- ovary (FTO) which comprises tumors located at ovaries or the adnexal mass. In total, we extracted DNA and RNA from 64 samples from 34 HGSC patients. For 28 patients, we had at least two tumor samples from different anatomical regions, while we had just one, either omental or tube-ovarian sample, for the remaining 7 patients. Informed consent was received from all the participating patients and the study was approved by the ethics committee of the Hospital District of Southwest Finland.

### DNA and RNA extraction

Qiagen AllPrep kit was used to extract DNA and RNA simultaneously from approximately 10 mg of tissue or 10^6^ cells which were then stored at -20° C and -80° C respectively. DNA concentration was measured using Quantus (Promega). To check its quality and integrity, 100 ng of the extracted genomic DNA was run on 1% agarose gel by gel electrophoresis.

### RNA-sequencing data analysis

We utilized RNA sequencing data generated from 42 tumor samples from 27 HGSC patients and 4 normal fallopian tubes (NFT) samples from unrelated individuals as a part of a separate study (https://www.deciderproject.eu/) to assess L1 transcriptional expression. Since L1 transcripts with 3’
s transductions can be detected with high reliability (Deininger et al., 2017; Philippe et al., 2016), we aimed to detect them using the bulk RNA sequencing data. Alignment files obtained in BAM file format were filtered for: i) all the reads that mapped in exonic regions and ii) all the reads that mapped to repeat regions by 90% using bedtools intersect (Quinlan and Hall, 2010). Forward reads and reverse reads were then separated using samtools (Danecek et al., 2021) using flags 67 and 131 respectively. These forward and reverse reads were then intersected with a 1kb region both upstream and downstream of all retrotransposition-competent (RC) L1s (Deininger et al., 2017) in a strand-aware manner. The RNA fragments mapping both upstream and downstream regions were counted using htseq-count (Anders et al., 2015). An RC-L1 locus was deemed to be “expressed” if the number of RNA fragments mapped on its 1 kb downstream window was at least 5 times as much as those mapped on its 1 kb upstream window.

### Differential Gene Expression and Gene-Set Enrichment Analysis

We processed, quantified, and deconvoluted the RNA sequencing reads as we previously described (Häkkinen et al., 2021). Briefly, read pairs were trimmed using Trimmomatic (version 0.33) (Bolger et al., 2014) and the trimmed reads were aligned to the GRCh38.d1.vd1 reference genome with GENCODE v25 annotation using STAR (version 2.5.2b) (Dobin et al., 2013). We quantified the gene level effective counts using eXpress (version 1.5.1-linux_x86_64)(Roberts and Pachter, 2013) and estimated the cancer cell-specific expressions using PRISM (Häkkinen et al., 2021).

We performed differential gene expression analysis between L1-high and L1-low samples on cancer cell-specific effective counts using DESeq2 (version 1.36.0) (Love et al., 2014). Gene set enrichment analysis (GSEA) (Subramanian et al., 2005) was performed using fgsea (version 1.22.0) (Korotkevich et al., 2021) by ranking differentially expressed genes based on their −log10(p-value)*sign(fold-change). We used the hallmark, KEGG, PID, and REACTOME gene sets collected from the Molecular Signatures Database (MSigDB, version 7.2) in GSEA.

### Genotyping PCR

To detect the presence/absence of two RC-L1s in their native genomic location, PCR primers were designed across the RC-L1 (**Figure 1A**). 1.5 ng of genomic DNA was amplified by conventional PCR using Phusion Hot Start II DNA polymerase (Thermo Scientific, catalog no.: F549L). The PCR mix was made as per the manufacturer’s instruction using the Phusion GC buffer to which 1.5 ng genomic DNA was introduced. PCR primers used and cycling conditions are provided in **Supplementary Table 4**. PCR products were then resolved on a 1% agarose gel.

### Reverse-transcriptase PCR

RT-PCR was done to detect the expression of RC-L1 mRNA. RT-PCR primers were designed such that the 3’
s-end of the forward primer specifically recognized RC-L1 specific ACA variant (Rodić et al., 2015) and the reverse primer annealed to the 3′ flanking non-repetitive region of the RC-L1 (**Figure 1A**). 1µg of RNA extracted from tumor samples and cancer cell lines was used to synthesize complementary DNA using SuperScript™ IV Reverse Transcriptase (Thermo Scientific, catalog no.: 18090010) using the manufacturer’s direction. Oligo dT primers (Thermo Scientific, catalog no.: 18418020) were used to ensure that mRNA with polyA tails are captured. cDNA produced was diluted in nuclease-free water at a 1:10 ratio. 1µl of diluted cDNA was then used for the RT-PCR reaction to detect L1-5219 and L1-5236 transcripts. Phusion Hot Start II DNA polymerase (Thermo Scientific, catalog no.: F549L) was used along with the Phusion HF buffer provided. The cycling conditions for RT-PCR and primers used are listed in **Supplementary Table 5**.

### Long-distance inverse PCR

LDI-PCR to detect somatic L1 retrotransposition was performed as previously described (Pradhan and Kauppi, 2019; Pradhan et al., 2017). Briefly, 100 ng genomic DNA extracted from tumor samples and cancer cell lines were digested using restriction enzymes *Nsi*I (New England Biolabs), *Vsp*I (FastDigest, Thermo Scientific), *Sac*I (FastDigest, Thermo Scientific), *Pst*I (New England Biolabs), *Hind*III (New England Biolabs) and *Spe*I (New England Biolabs) according to manufacturers’ instruction. Restriction enzymes were heat-inactivated after digestion reaction, and digested DNA was self-ligated to form a circular DNA template using T4 DNA ligase (Thermo Scientific, catalog no.:EL0011). The standard reaction mix was prepared as per the manufacturer’s instruction except for *Vsp*I digested DNA, to which Polyethylene glycol (PEG) was added to a final concentration of 5% w/v. The ligase reaction mix was directly added to the digested DNA and incubated at room temperature for 1 hour, followed by heat inactivation of the enzyme. *Vsp*I-digested DNA was exceptionally incubated in ligase mix at 4° C overnight and heat-inactivated subsequently. Finally, touch-down PCR was performed on thus generated circular DNA at optimized annealing temperature (for cycling conditions, see **Supplementary Table 6**). Primers used to detect somatic retrotranspositions arising from L1-5219 and L1-5236 and their optimized annealing temperature are listed in **Supplementary Table 6**. Circular DNA template libraries generated by *Nsi*I, *Vsp*I, *Sac*I, & *Pst*I, and *Nsi*I, *Vsp*I, *Hind*III & *Spe*I was used for L1-5219 and L1-5236 mediated insertion detection, respectively.

### Nanopore sequencing

In total, 20 different LDI-PCRs were performed per sample (12 reactions for L1-5219 using circular templates generated from 4 different restriction enzymes and amplifying each with 3 sets of primer pairs; and 8 reactions for L1-5236 using circular templates generated from 4 different restriction enzymes and amplifying each with 2 sets of primer pairs), with two technical duplicates for each reaction. All LDI-PCR products from each sample were pooled and quantified using Quantus (Promega). 5-20 ng of the pooled PCR product was analyzed using Tapestation (Agilent), and the average molecular weight of the pooled amplicon was calculated. 0.1 pmol of the pooled PCR product from each sample was bar-coded using ligation-based barcoding kit NBD- 104 and NBD-114 (Oxford Nanopore Technologies). Barcoded amplicon libraries were pooled in an equimolar fashion in order to produce the final sequencing library which was introduced to a FLO-MIN106D flowcell and run on MinION Mk1B. The raw fast5 files were base-called using the high-accuracy base-calling option on Guppy Software version 3.4.5 (Oxford Nanopore Technologies).

### LDI-PCR/Nanopore-seq data analysis

In order to identify the target genomic location of somatic L1 retrotransposition, we utilized a structural variation (SV) calling pipeline called pipeline-structural-variation (available at https://github.com/nanoporetech/pipeline-structural-variation). In short, the pipeline employs minimap2 (Li, 2018) to map the fastq sequencing reads to the human reference genome, calculates suitable parameters based on the read depth (Pedersen and Quinlan, 2018), and finally calls the structural variation using Sniffles (Sedlazeck et al., 2018). The minimum read length was set to 200 bp, and the rest of the parameters were kept to default. Since Sniffles is designed to detect SV from whole-genome sequencing data, it erroneously detects the restriction enzyme cut-site generated in our LDI-PCR protocol as breakpoints. We therefore manually curated the SV calls generated, by visualizing them on the Integrated Genome Viewer (IGV) and by analyzing the reads associated with each call. In order to qualify as “true” SV calls, the reads associated with SV detected had to meet the following criteria: a) be disjointed at one of the restriction enzyme cut-sites and b) have supplementary alignments to the unique tag of L1-5219 or L1-5236. Additionally, we visualized the alignment files for each sample at the breakpoints to observe polyA sequence and target-site modification (duplications or deletions) that are typical sequence signatures of L1 retrotransposition and reported them in **Supplementary Table 1**.

### DNA-sequencing data analysis

We employed 24 WGS tumor samples from 24 patients together with their matched normals from the DECIDER project (https://www.deciderproject.eu/) to analyze the genomic features. Briefly, raw reads were trimmed using Trimmomatic 0.32 (Bolger et al., 2014) before alignment to GRCh38.d1.vd1 using BWA-MEM 0.7.12-41039 (Li, 2013). This was followed duplicate marking with Picard 2.6 (https://github.com/broadinstitute/picard) and base quality recalibration with GATK 3.7 (McKenna et al., 2010). We called somatic variants with GATK 4.1.9.0 Mutect2 with joint calling and germline variants with GATK with allele-specific filtering and joint-genotyping, both according to best practices. Germline status in tumor samples was assessed via forced calling with Mutect2. Allele-specific copy-numbers were estimated using ASCAT (Van Loo et al., 2010) with segmentation from GATK. We included only samples with at least 10% purity.

We computed SBS and ID spectra according to mutation classification of COSMIC reference signatures v3.2 (Tate et al., 2019). SBS signatures were adjusted for GRCh38 trinucleotides sans chromosomes Y and M. We fitted SBS and ID signatures independently on each sample using an R implementation based on SigProfilerAttribution (Alexandrov et al., 2020). Signatures with at least 20% ovarian cancer occurrence in COSMIC were selected as common signatures.

### Detection of inactivating variant in L1-5219 sequence

To detect the presence inactivating alleles in the L1-5219 sequence, we compared the variants present in the L1-5219 sequence in the genomes of L1-5219^**-**^ samples (samples with no somatic L1 insertion arising from L1-5219) with that of L1-5219^**+**^ samples. We applied LONGSHOT (Edge and Bansal, 2019), a tool to detect single nucleotide variants (SNV) in single-molecule long-read sequencing data, to identify SNVs in L1-5219 and its flanking sequence. BCFtools (Li, 2011) was then used to identify L1-5219 variants specific to L1-5219^**-**^ samples.

### Data visualization

All the plots presented in this paper were created in R (https://www.R-project.org/) using ggplot2 (Wickham, n.d.), genomicRanges (Lawrence et al., 2013) and Rcircos (Zhang et al., 2013) packages.

## Acknowledgements

We would like to thank Dr. Manuela Tumiati and Taina Turunen for technical assistance and sample collection. We would also like to acknowledge CSC – IT Center for Science, Finland, for computational resources and Biomedicum Functional Genomics Unit (FuGu) for genomics services.

