## Supplementary Material for "L1 retrotransposition is regulated post-transcriptionally In High-Grade Serous Ovarian Cancer"

### Supplementary Results

#### *1. Characteristics of somatic L1 insertions arising from two RC-L1*

Long reads generated by LDI-PCR/Nanopore-Seq allowed the identification of 5´ and 3´ junctions of the somatic L1 insertions, along with the entire inserted sequence in one single read. We used this nucleotide-level information from representative reads of each of the 233 somatic L1 insertions to characterize them in greater detail (**Supplementary Table 1**).

The L1 retrotransposition process initiates with the formation of a nick in a T-rich region of the genome which serves as an annealing partner for the polyA tail of L1 mRNA. This T-rich region in the target locus is then used as a primer for reverse-transcription of the L1 mRNA in a process referred to as Target-Primed Reverse Transcription (TPRT) (Luan et al., 1993), which leads to polyA sequence on 3´ end of somatic L1 insertions. We observed this 3´ polyA sequence in 82% of *de novo* L1 insertions. L1 retrotransposition also often leads to duplication or deletion of the target genomic location (Gilbert et al., 2002). While target-site deletion of 1-776 bp (Median = 5 bp) was observed in 20 L1 insertions, target-site duplication of 1-220 bp (Median = 14 bp) was much more common, observed in 206 L1 insertions. The remaining 7 insertions occurred without any loss or duplication of target-site DNA sequence, suggesting that both first as well as second strand nicks were made precisely at the same genomic coordinate. 5´ inversion of the L1 inserts by twin-priming (Ostertag and Kazazian, 2001) was observed in 34% and 26% of L1 insertions stemming from L1-5219 and L1-5236, respectively.

Since our assay detects L1 retrotransposition events that involve mobilization of non-repetitive sequence on the 3´ flank by 3´ transduction, we had sequence information of the 3´ transduced region for all of the somatic L1 insertions detected. As 3´ transduction occurs due to the usage of an alternative polyadenylation signal (PAS) downstream of L1's own canonical PAS to terminate transcription, we could estimate alternative PAS usage in all 233 somatic L1 insertions. For L1-5219, 6th and 9th PAS at chr22:28669887 and chr22:28670116 were the most preferred transcription termination signals (**Figure 3D**). In the case of L1-5236, the 4th PAS at chrX:11714563 was the preferred transcription termination signal (**Figure 3D**). However, it should be noted that due to primer design constraints, we were unable to detect L1-5219 transductions that utilized the 2nd PAS, and L1-5236 transductions that utilized 2nd and 3rd PAS.

All somatic L1 insertions detected were truncated towards the 5´ end, thus none of the somatic L1 insertions identified in our HGSC sample cohort could possibly yield new L1 retrotransposition events. 64% of the somatic L1 insertions were truncated to the extent that they did not contain any repetitive L1 sequence but instead retained only the 3´ transduced element in the insertion site (**Figure 3D**). Such insertion events are often referred to as orphan transductions.

To summarize, we uncovered different L1 retrotransposition hallmarks, such as 3´ polyA sequence, target-site modification, and 5´ inversion by analyzing LDI-PCR/Nanopore-Seq generated sequencing reads. We identified the most preferred PAS on the 3´ flanking genomic region of L1-5219 and L1-5236 that were used for transcription termination and hence to generate a 3´ transduced region. Moreover, our data demonstrate that orphan transductions are remarkably frequent in HGSC.

#### *2. Somatic L1 insertions are concentrated in less-expressed genes.*

As somatic L1 insertions in the intronic region of the gene can cause alteration in gene expression (Nguyen et al., 2018; Shukla et al., 2013), we investigated the intragenic somatic L1 insertions and their effect on the expression of the target genes. 115 out of 233 somatic L1 insertions found occurred in the intron of a protein-coding or long non-coding RNA gene. We checked the expression of these host genes in 42 out of 66 tumor samples used in this study for which we had RNA-seq data and found that 109 of them were expressed in very low levels (median TPM < 1). Out of 7 genes with a median of more than 1 TPM in the tumor samples, that harbored a somatic L1 insertion, we could only assess the impact of L1 insertion in gene expression for two genes (*CDK13* and *SYNE2*) as RNA-seq data was not available for all the HGSC tumor samples. While *CDK13* was upregulated in the tumor sample that showed somatic L1 insertion in it (TPM = 7.5 vs median TPM of 4.3 in 42 tumor samples), *SYNE2* was slightly downregulated (TPM = 11.9 vs median TPM of 15.6 in 42 tumor samples). The inserted L1 sequence in both the genes was in the same orientation as the gene (referred to as sense orientation in **Supplementary figure 5**). Both somatic L1 insertions in *CDK13* and *SYNE2* were, however, detected by 6 and 44 LDI-PCR/Nanopore-Seq reads in the omental tumor of the EOC781 patient respectively and thus can be subclonal. For reference, other 14 insertions identified in the same tumors were supported by 8-679 reads (Median=160) after normalizing for the number of circular DNA templates that could detect each insertion. Therefore, it is possible that the expression changes due to somatic L1 insertions in these genes can be masked by other tumor cells in the bulk tumor mass that have not acquired the insertion.

#### *3. Somatic L1 insertions exhibit intratumor heterogeneity*

Different somatic L1 insertions detected in the same tumor sample were represented by varying numbers of Nanopore reads generated after sequencing LDI-PCR products which suggested the subclonal nature of these insertions. Although subclonal insertions can be inferred by these varying number of reads, there are several technical characteristics of the LDI-PCR method that needs to be considered. First, the size of the circular DNA template containing a somatic L1 insertion produced as the result of self-ligation following restriction enzyme digestion dictates how efficiently they are amplified during the LDI-PCR process and thus has a direct impact on the number of reads produced (**Supplementary figure 6**). Second, the number of reads supporting a somatic L1 insertion is also affected by the number of restriction enzymes that are able to make a PCR-amplifiable circular DNA template containing the insertion in question. Third, the number of reads is also affected by the availability of primer-binding sites in the 3´ transduced genomic region.

Thus for simplicity, we only selected those L1 insertions that were detected from a circular DNA template generated by just one restriction enzyme and classified them according to the primer pair(s) that could amplify it in two HGSC samples exhibiting the highest number of somatic L1 retrotransposition (EOC781 Ova and EOC124 Ova(R), **Supplementary figure 7**). While most of the insertions followed the known inverse relationship between the circular DNA template size, which can be inferred by the Nanopore sequencing read length of the LDI-PCR product, and the total number of Nanopore sequencing reads within the primer-specific group, there were some that deviated from this pattern (**Supplementary figure 7**). Moreover, some insertions, despite being amplified by two or even three different primer pairs in two and three respective PCR reactions were detected by fewer reads than other insertions of similar read length that were detected by just one primer pair (**Supplementary figure 7**). These results indicate the subclonal nature of somatic L1 insertions within a tumor sample.

##### *4. Clinical features of HGSC patients with different L1 status*

We asked whether L1 insertion status of HGSC patients associates with their clinical features. To do so, we chose for each patient the tumor with highest somatic L1 insertions, whenever there was more than one tumor sample from the same patient. In cases where we had only one tumor sample from a patient, only those with L1-high samples were considered to avoid false categorization of a patient as L1-null or L1-low. 29 out of 35 patients qualified for this criterion.

Since we observed a positive correlation between the total number of somatic L1 retrotransposition events in a tumor and the mutational signature that were associated with age of the cancer patient, we hypothesized that the age of the patient at diagnosis may correlate positively with the number of L1 insertions observed in their tumors. However, we did not find any correlation between the age versus total somatic L1 insertion acquired (**Supplementary Figure 8A**). The average age of the patients in the L1-high, L1-low and L1-null group was 67, 68 and 64 respectively. When we considered all the patients that exhibited at least one somatic L1 insertion in one group of L1-positive patients, their average age at diagnosis was 67 compared to 66 in the L1-null group. Somatic L1 insertion status was not associated with primary therapy outcome, although all four cases with highest L1 burden were in the complete response group (**Supplementary Figure 8B**). Furthermore, there was no statistically significant enrichment of patients with certain cancer stages or those within a particular treatment arm (Primary Debulking Surgery (PDS) or Neoadjuvant Chemotherapy (NACT)) in L1-null patients versus L1-positive patients (**Supplementary Figure 8C-D)**.

#### Supplementary Figures

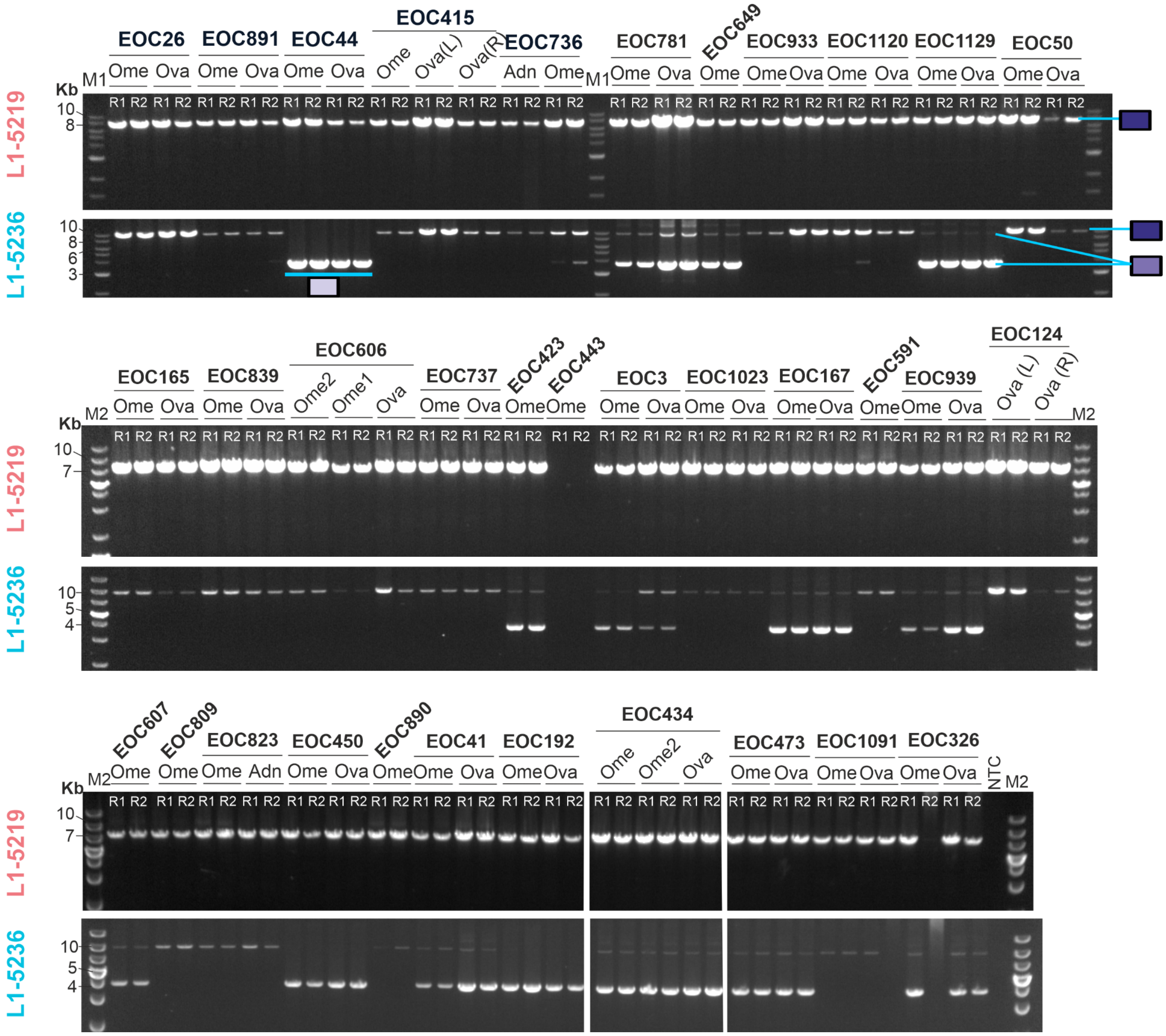


**Supplementary Figure 1**: **Determination of presence/absence of RC-L1s by genotyping PCR assay.** Agarose gel image showing the result of the genotyping PCR for two RC-L1s, L1-5219 and L1-5236, assayed in 64 HGSC tumor samples included in this study. PCR primers were designed across the full-length L1 sequence. As a result, a PCR product of 7.2 and 9.8 kilobases (Kb) is observed when L1-5219 and L1-5236 were present in the genome respectively. When the RC-L1s are absent, a PCR product 6 Kb shorter than the aforementioned amplicon is observed. Three types of cases are labeled on the top gel image –

dark purple= RC-L1 present in both homologous chromosomes (homozygous present); purple = RC-L1 present in one copy only (heterozygous present) and pale purple = both homologous chromosomes lack the RC-L1. Genotyping PCR failed in one sample- EOC443 Ome. (R1 and R2 are two replicates of the same reaction. M1 = 1kb DNA ladder (New England Biolabs); M2 = 1 kb plus DNA ladder (ThermoScientific))

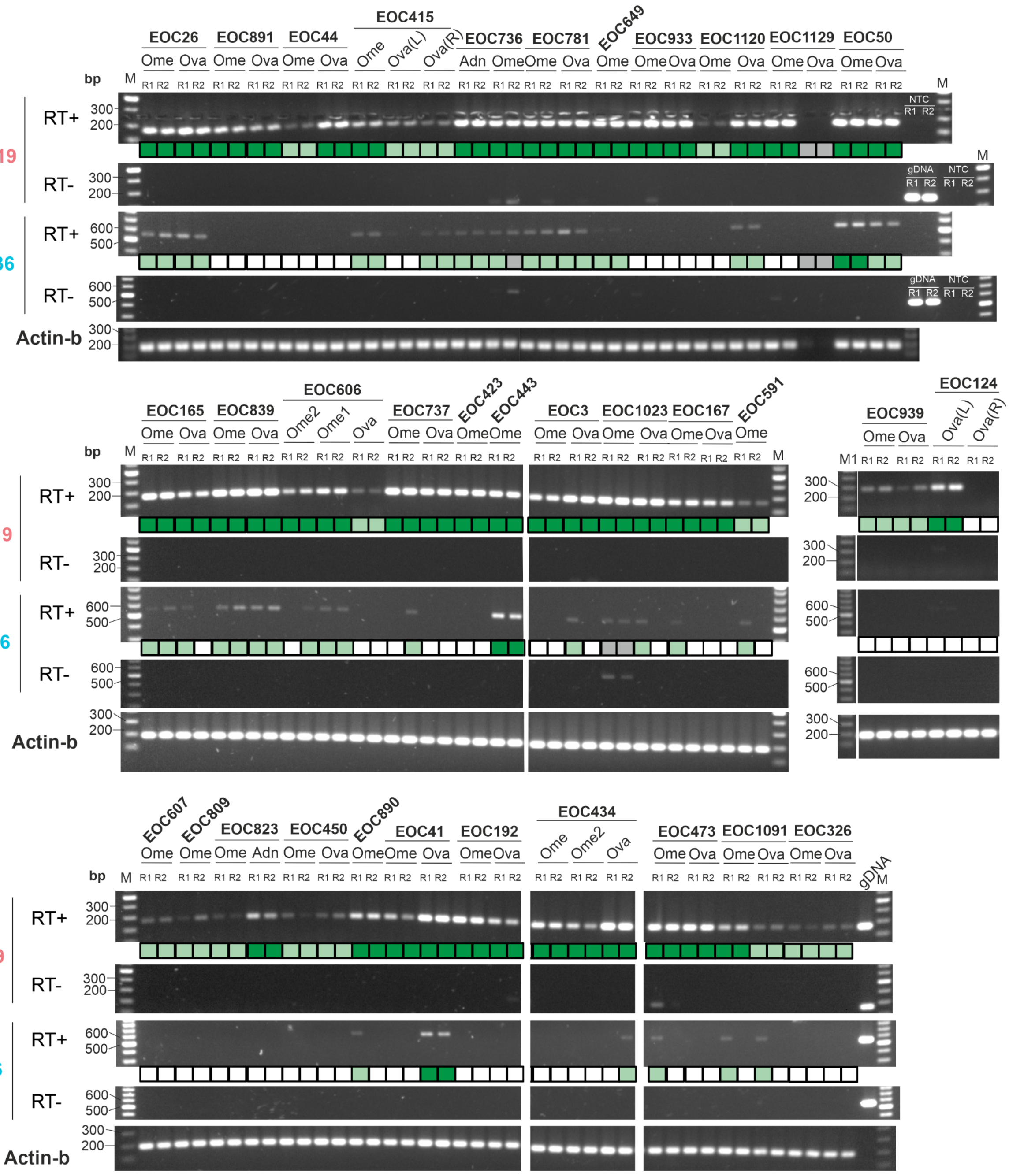

**Supplementary Figure 2**: **RNA expression of L1-5219 and L1-5236 by RT-PCR**. Agarose gel image showing result of RT-PCR which was done to assess the transcriptional activity of two RC-L1s – L1-5219 and L1-5236 in 66 HGSC tumor samples included in this study. Semi-quantitative scoring of RC-L1 expression was done and labeled by dark-green, light-green and white boxes representing strong, weak and no expression respectively. Expression of the *ACTB* gene was used as a reference for RNA integrity. RT-PCR product size for L1-5219, L1-5236 and *ACTB* are 262, 547 and 200 bp respectively. (R1 and R2 are two replicates of the same reaction. M = 100bp DNA ladder (ThermoScientific); M1 = 100 bp plus DNA ladder (ThermoScientific))

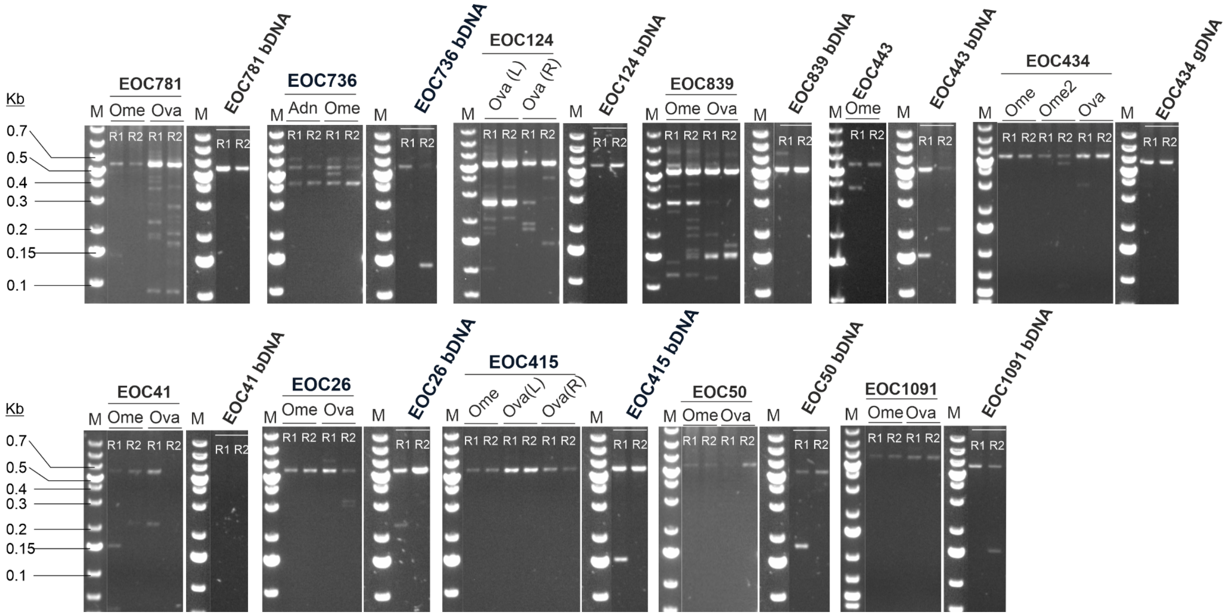


**Supplementary Figure 3**: **L1 insertions detected by LDI-PCR are tumor-specific.** Side-by-side comparison of agarose gel displaying LDI-PCR results obtained from tumor samples from L1-high patient (patients having at least one L1-high tumor sample) with genomic DNA extracted from their peripheral blood sample (labeled as “bDNA”). LDI-PCR was done using L1-5219 specific inverse primers on *Vsp*I generated circular DNA template. Native RC-L1 specific LDI-PCR product generated by this reaction is 5.6 kb in length and is visible in tumor as well as normal sample (EOC41 bDNA was an exception where the PCR failed). Reproducible LDI-PCR product of sizes different than that of the native product was only observed in tumor samples and not in blood samples. (R1 and R2 are two replicates of the same reaction. M = 1 kb plus DNA ladder (ThermoScientific))

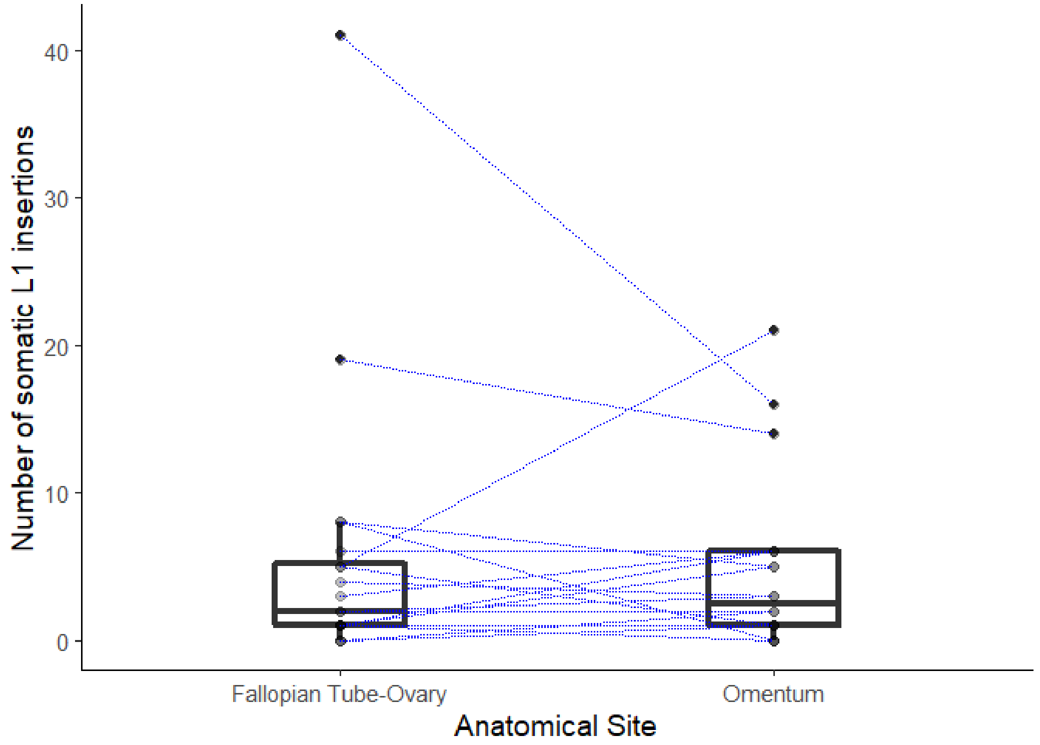


**Supplementary Figure 4**: **Pair-wise comparison of somatic L1-insertion frequency in tumors found in Fallopian Tube-Ovarian (FTO) vs. Omentum from the same patient.** In cases with more than one tumor of Omental or FTO origin from the same patient, tumors with the highest number of somatic L1-insertions were selected.

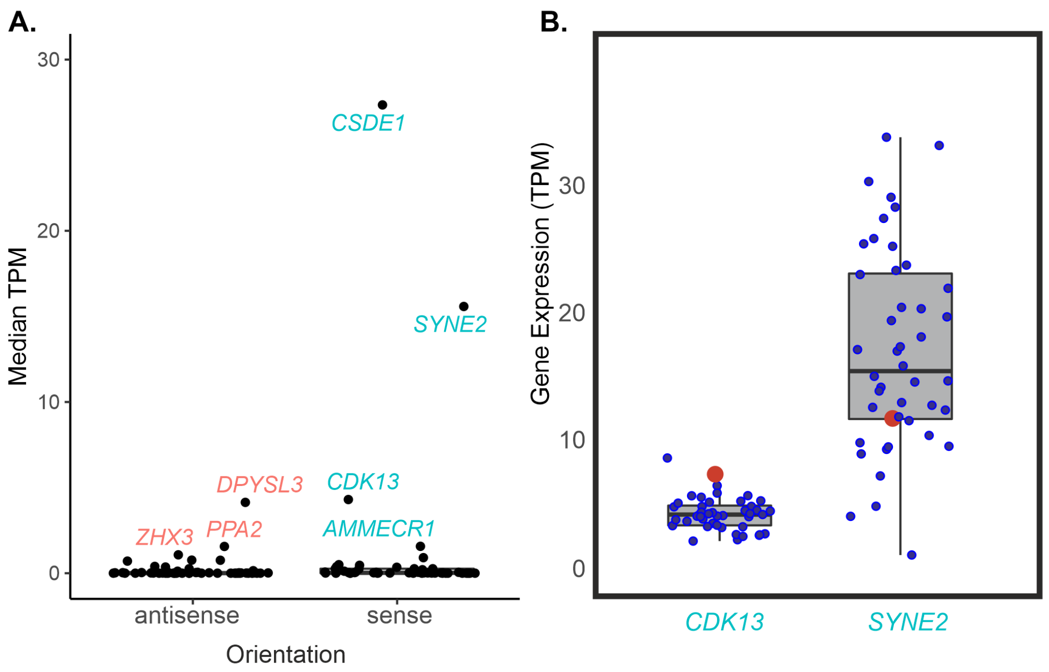


**Supplementary Figure 5. RNA expression of L1 target genes**. A) Median expression values (in TPM) of genes that harbored L1 insertion in one tumor sample across 40 HGSC tumor samples. L1 target genes that harbor somatic L1 insertions in the same orientation as theirs are classified as "sense" and those that have insertions in opposite direction are labeled as "antisense". All the genes that have median TPM>1 are labeled by their respective gene names. B) RNA expression of the gene that harbored a somatic L1 insertion in a tumor sample (red circle) is shown in the context of its expression in other tumor samples (blue circles) in the cohort.

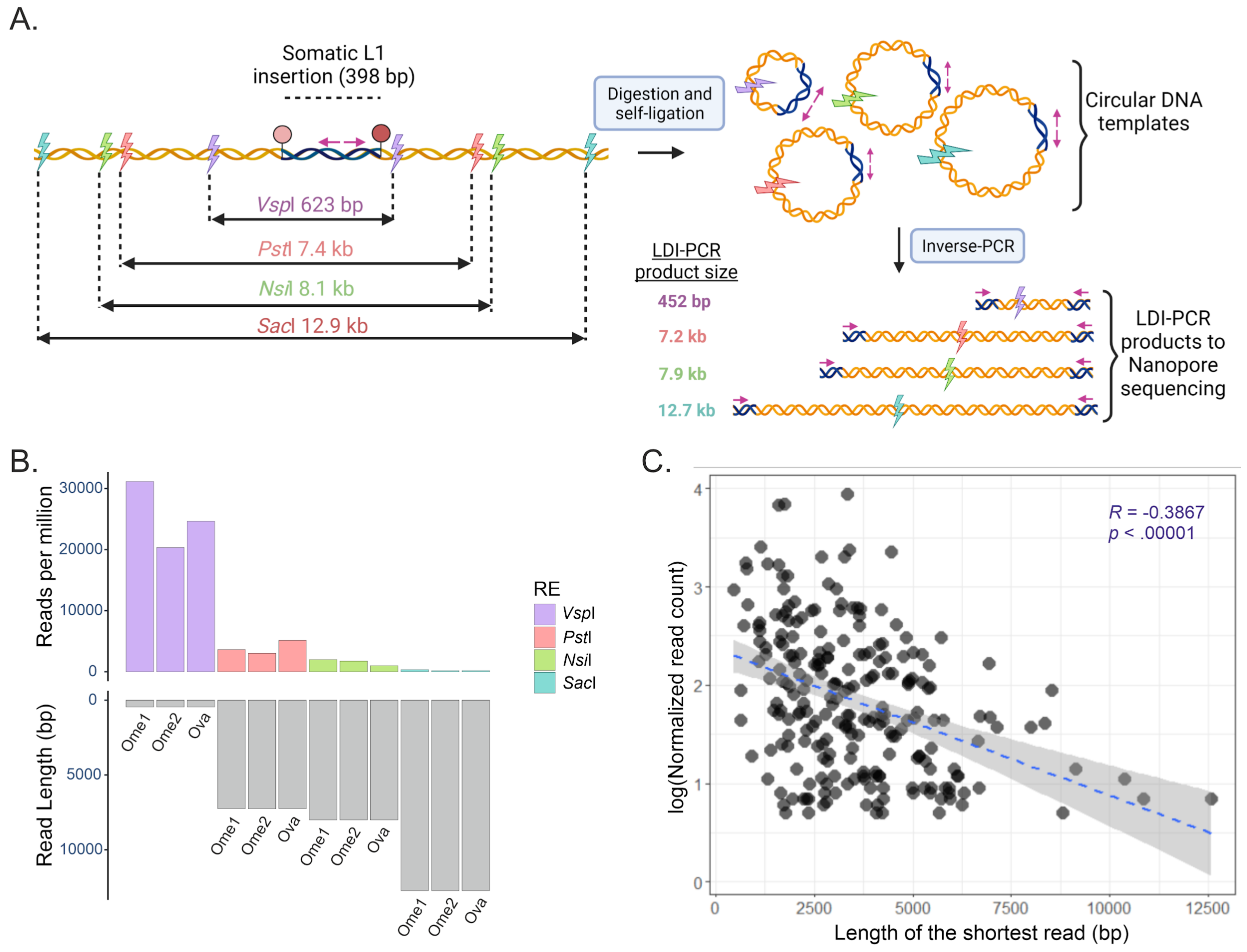


**Supplementary figure 6**: **Relationship between circular DNA templates of different sizes generated by different restriction enzymes as inferred by LDI-PCR/Nanopore read length**. A) Schematic representation of an orphan L1 insertion detected in three tumor samples (two omental tumors labeled Ome1 and Ome2 and one ovarian tumor labeled as Ova in B) from one HGSC patient. This insertion originated from L1-5219 and was detected from circular templates generated by all four restriction enzymes used. B) LDI-PCR/Nanopore-Seq reads generated from circular templates created by each restriction enzymes were normalized by total reads generated from each sample and plotted along with its respective read length to show the inverse relationship between the reads generated and the length of the read. C) As the number of reads decreased in an exponential fashion corresponding to the read length and the shortest read contributed to most of the read counts, we plotted the log of Normalized read count with the length of shortest read (when an insertion was detected from template generated by different restriction enzymes leading to different read lengths) for all 233 somatic L1 insertions detected in this study. Normalized read count = total reads supporting an L1 insertion/number of restriction enzymes detecting it.

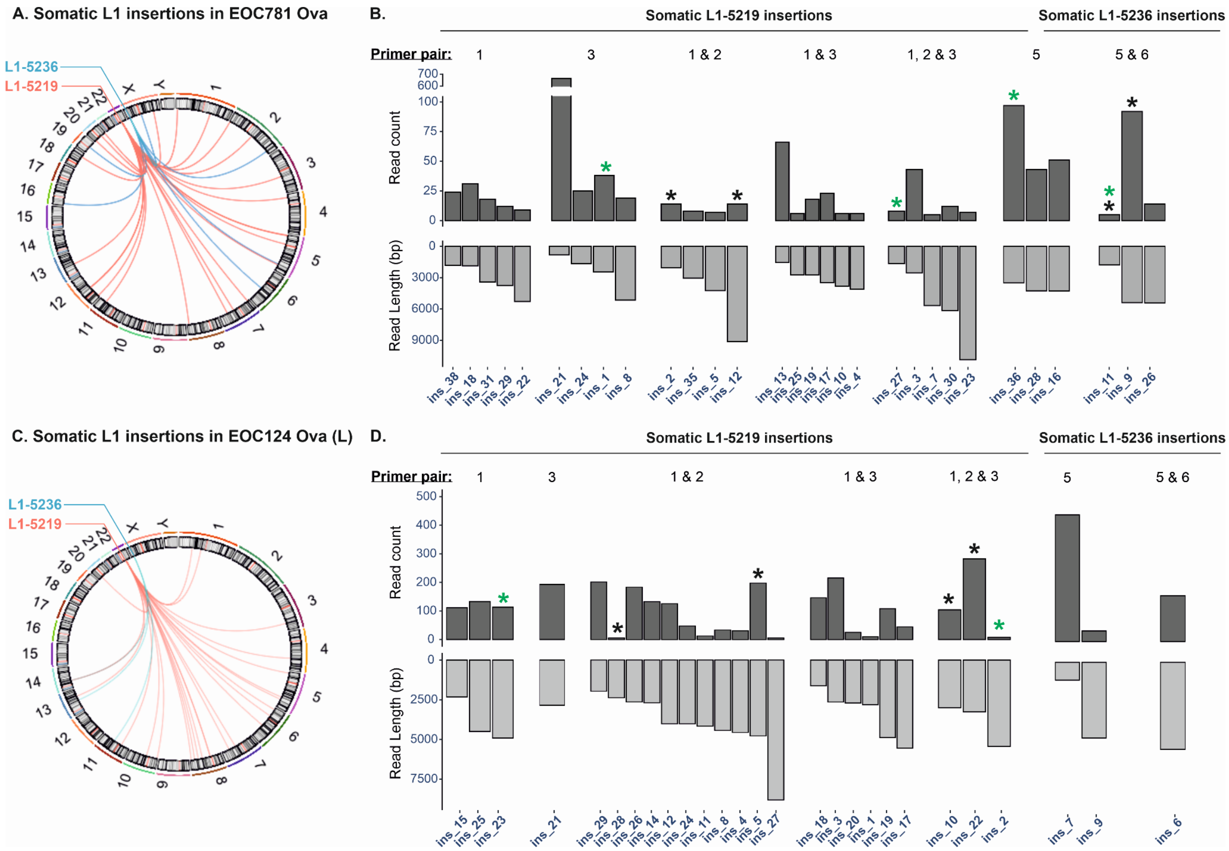


**Supplementary Figure 7**: **Subclonal nature of somatic L1 retrotransposition events.** The total number of Nanopore reads detecting each somatic L1 insertion in tumor samples (EOC781 OvaL (A) and EOC124 OvaR (B)) is plotted alongside their corresponding read length. All the somatic L1 insertions represented were detected by circular DNA templates generated by a single restriction enzyme. Insertions detected by different primer pair(s) are grouped accordingly. L1 insertions that do not follow the inverse relationship between the read length and read count are marked with a green asterisk. Black asterisk labels L1 insertion pairs of comparable read length that are detected by one vs. more than one PCR primer pair.

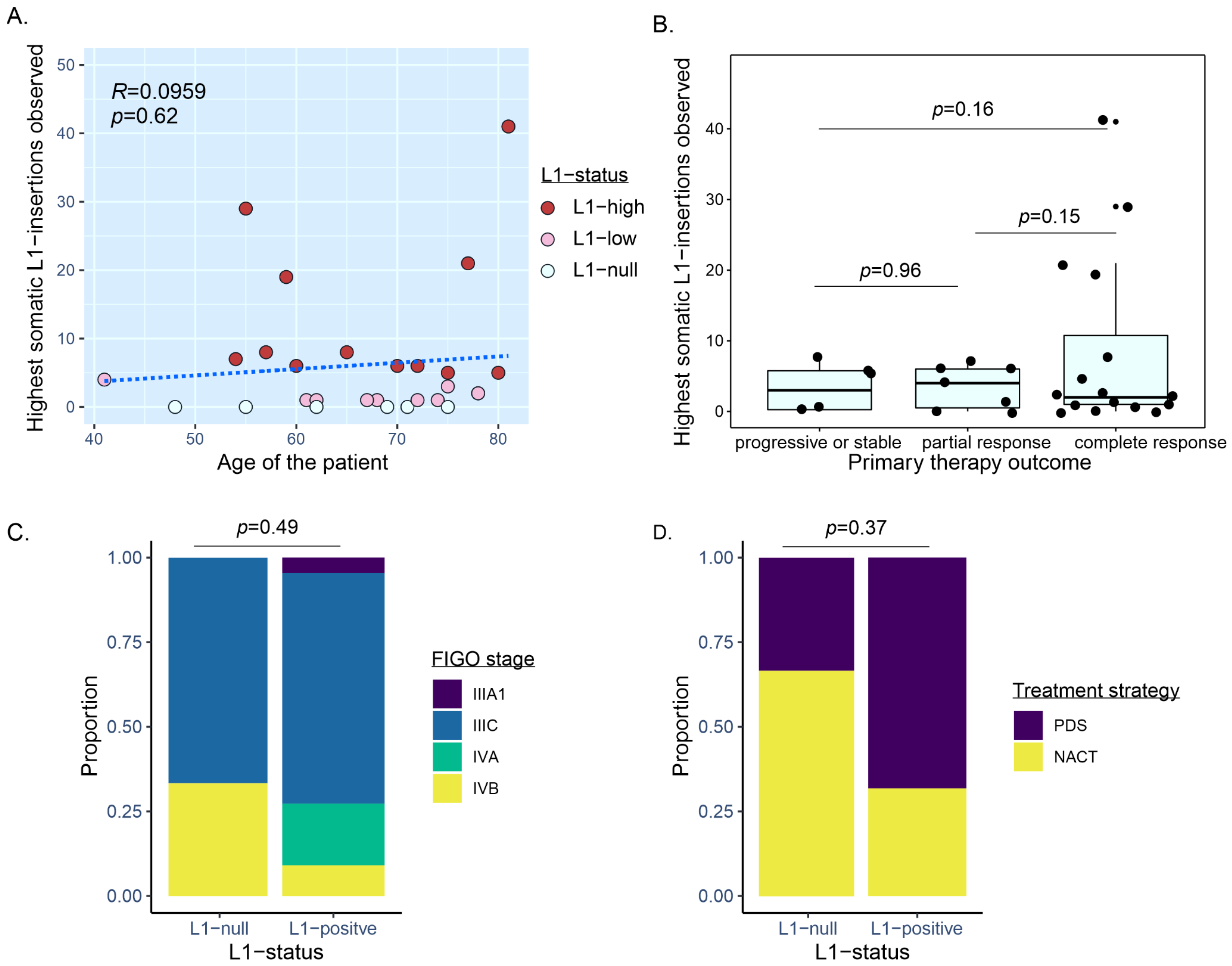


**Supplementary Figure 8**: **Relationship between somatic L1 retrotransposition activity of HGSC patients and their clinical features** such as age at diagnosis (A), the primary therapy outcome of the disease (B) Federation of Gynecology and Obstetrics (FIGO) stage of disease (C), the treatment strategy used (D). The highest number of somatic L1-insertion observed in a patient tumor is compared with age and primary therapy outcome (A and B respectively, p-value obtained after a two-tailed student's t-test is indicated). To observe the association between L1 retrotransposition activity and clinical stage (C) and treatment strategy (D) patients are classified as L1-null (with no somatic L1 insertions, n=6) and L1-positive (with at least 1 somatic L1 insertion, n=22). P-values were obtained as a result of Fisher's exact test.

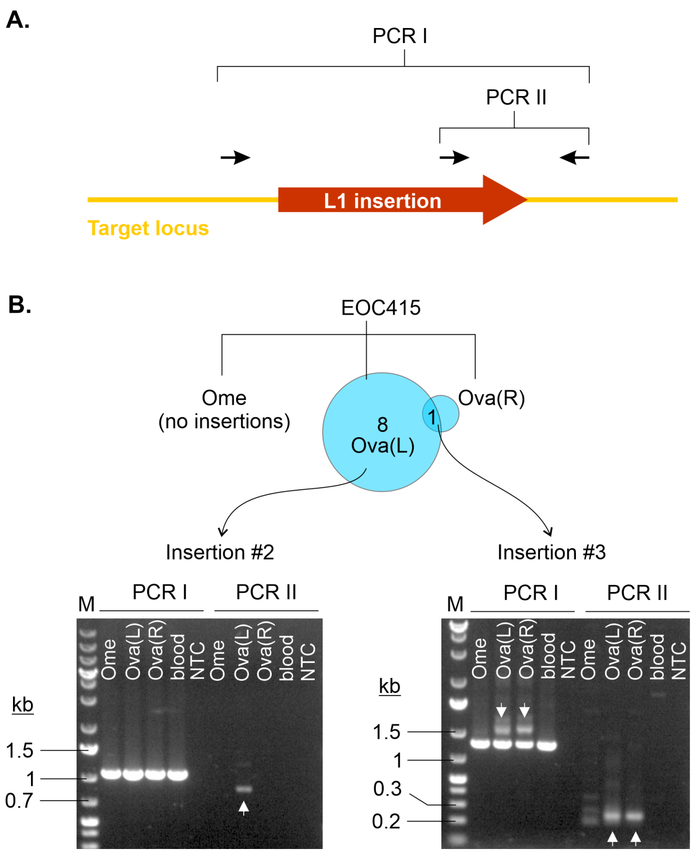


**Supplementary Figure 9**: **Validation of the somatic L1 insertion mediated intrapatient heterogeneity detected by LDI-PCR/Nanopore-Seq.** A) PCR assays (PCR I and II) validating somatic L1 insertions are shown with primers symbolized by black arrows. B) Validation PCRs were done for somatic L1 insertions (insertion #2) uniquely found in tumor located at left ovary (Ova(L) and another (insertion #3) that was shared between tumors located at left and right ovary (Ova(R)). PCRII shows that insertion #2 is in fact uniquely present in Ova(L) tumor. Both PCR I and II shows that insertion #3 is present in Ova(L) as well as Ova(R) but absent in Ome as well as normal genomic DNA extracted from the patient's blood sample (lane labeled as "blood"). Affirmative PCR products of correct size indicating the presence of somatic L1 insertions are pointed by white arrows. (M = 1 kb plus DNA ladder (ThermoScientific); NTC= No Template Control)

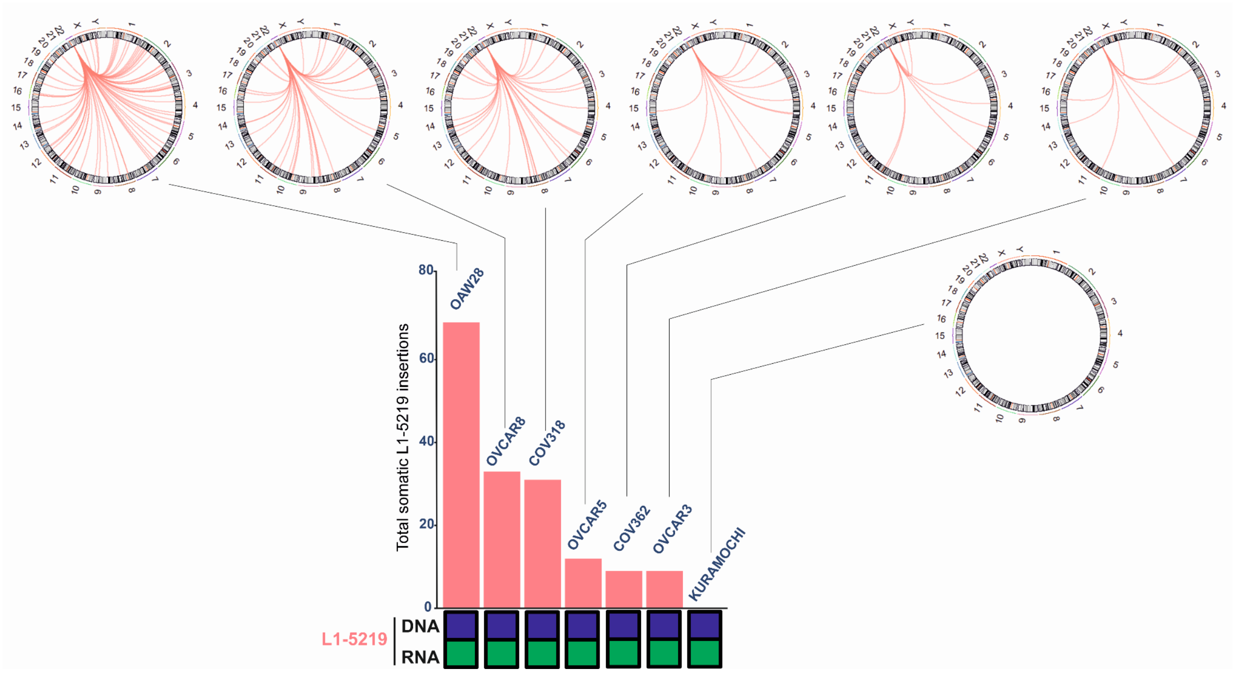


**Supplementary Figure 10. Retrotransposition activity of L1-5219 in commercial HGSC cell lines.** Frequency of somatic retrotransposition stemming from L1-5219 in 7 HGSC cell lines as detected by LDI-PCR/Nanopore-Seq. L1-5219 was homozygously present and transcribed in all cell lines (symbolized by dark purple and dark green boxes respectively. Circos plots show patterns of retrotransposition originating from L1-5219 located at chromosome 22.

Shukla R, Upton KR, Muñoz-Lopez M, Gerhardt DJ, Fisher ME, Nguyen T, Brennan PM, Baillie JK, Collino A, Ghisletti S, Sinha S, Iannelli F, Radaelli E, Dos Santos A, Rapoud D, Guettier

C, Samuel D, Natoli G, Carninci P, Ciccarelli FD, Garcia-Perez JL, Faivre J, Faulkner GJ. 2013. Endogenous retrotransposition activates oncogenic pathways in hepatocellular carcinoma. *Cell* **153**:101–111.
